# Targeted cortical stimulation reveals principles of cortical contextual interactions

**DOI:** 10.1101/2022.06.22.497254

**Authors:** Shen Wang, Agostina Palmigiano, Kenneth D. Miller, Stephen D. Van Hooser

## Abstract

Cross-orientation suppression is a classic form of contextual normalization in visual cortex. There is substantial evidence that feed-forward processes contribute to cross-orientation suppression, but binocular studies suggest there may also be a contribution from cortical circuits. We used a combined imaging/optogenetic stimulation system to provide patterned optogenetic stimulation to cells in orientation columns that either matched or were orthogonal to the preferred visual orientation of neurons recorded with electrodes. When visual or optogenetic stimulation of columns preferring one orientation was combined with optogenetic stimulation of columns preferring the orthogonal orientation, we observed less suppression than when orthogonal stimulation was provided visually, suggesting that cortical circuits do not provide a large fraction of visual cross-orientation suppression. Integration of visual and optogenetic signals was linear when neurons exhibited low firing rates and became sublinear when neurons exhibited higher firing rates. We probed the nature of sublinearities in cortex by examining the influence of optogenetic stimulation of cortical interneurons. We observed a range of responses, including evidence for paradoxical responses in which interneuron stimulation caused a decrease in inhibitory firing rate, presumably due to the withdrawal of recurrent excitation. These results are compatible with cortical circuits that exhibit strong recurrent excitation with stabilizing inhibition that provides normalization, albeit normalization that is too weak across columns to account for cross-orientation suppression.

**Significance statement:** When multiple stimuli are presented to the cortex simultaneously, the response is usually less than the sum of the responses individually, a phenomenon called normalization. Up to now it has been unclear whether and in what circumstances cortical circuits themselves contribute to cross-orientation normalization. Prior studies have found a lack of direct inhibition within the cortex that could underlie cross-orientation normalization, and other studies suggested that the entire phenomenon could be a property of the feed-forward input to the cortex. Consistent with this feed-forward hypothesis, we found that we could not induce cross-orientation suppression by direct stimulation of the cortex. We did find evidence for cortical normalization, but not cross-orientation normalization that is strong enough to account for cross-orientation suppression.

## 2. Introduction

In visual cortex, neurons respond to bars or gratings of a preferred orientation but are also highly influenced by the simultaneous presentation of additional stimuli, such as gratings that are orthogonal to the preferred orientation or flanking stimuli (Bishop et al., 1973; Morrone et al., 1982; Bonds, 1989; DeAngelis et al., 1992; Das and Gilbert, 1999; Cavanaugh et al., 2002; Smith et al., 2006; Busse et al., 2009; MacEvoy et al., 2009). Many of these contextual influences are well-described phenomenologically by a normalization equation that consists of summation and division (Bonds, 1989; Heeger, 1992; Carandini and Heeger, 2011). However, the cortical circuit mechanisms that underlie contextual processing are incompletely understood.

A classic form of normalization is cross-orientation suppression: the response of visual cortical neurons to a preferred orientation is suppressed when an orthogonal stimulus is also presented, even if the neuron exhibits no response to the orthogonal stimulus when presented alone. Cross-orientation inhibition emerged as an early hypothesis for the circuit implementation (DeAngelis et al., 1992; Heeger, 1992), but experiments that blocked cortical inhibition or measured the orientation dependence of cortical inhibition did not find evidence consistent with this hypothesis (Anderson et al., 2000; Katzner et al., 2011). Subsequently, careful consideration of the responses of the feed-forward inputs from the lateral geniculate nucleus (LGN) led to a feed-forward hypothesis: that desynchronization of the temporal response phases of the LGN inputs to a given V1 cell, LGN saturation and rectification, and V1 spike threshold could account, in large part, for the reduced responses observed in cross-orientation suppression (Freeman et al., 2002; Li et al., 2006; Priebe and Ferster, 2006). Recent experiments further showed that, in mouse, suppressing cortical spiking does not impact subthreshold normalization (Barbera 2022), suggesting cortical circuits did not contribute. However, experiments using dichoptic presentation of the two gratings showed a cortical component of cross-orientation suppression (Sengpiel and Vorobyov, 2005), implying a cortical contribution, and it remains unclear whether the arguments supporting the feed-forward hypothesis, which were based on analysis of responses to drifting gratings, would apply to the thin bar stimuli used by MacEvoy et al. (2009), which should cause less desynchronization of putative LGN inputs (Lauritzen et al., 2001). Experiments and circuit models suggest that recurrent cortical connections, particularly those that operate in an inhibition-stabilized regime (Ozeki et al., 2009; Shushruth et al., 2012; Rubin et al., 2015; Litwin-Kumar et al., 2016; Sato et al., 2016; Palmigiano et al., 2020), could implement normalization that does not involve direct cross-orientation inhibition.

We designed experiments to directly probe the role of cortical circuits in cross-orientation normalization. Using a custom ProjectorScope apparatus (Huang et al., 2014; Roy et al., 2016), we located orientation maps in ferret visual cortex with intrinsic signal imaging, and then provided patterned optogenetic stimulation directly to different orientation columns. We examined, with electrodes, how cortical circuits integrated visual and optogenetic stimuli of different contrasts and orientations.

Optogenetic activation of an orientation column caused spiking activity that spread beyond the orientation columns that were directly stimulated (but not when glutamatergic synapses were blocked). Paired responses to visual stimulation at the preferred orientation and optogenetic stimulation of the orthogonal columns were smaller than the linear sum of the two stimuli alone, but the paired suppression was substantially less than was observed with purely visual paired stimuli, suggesting that cross-column suppression is much too weak to explain cross-orientation suppression. Responses differed depending upon whether stimulation was delivered visually, optogenetically, or mixed, which indicated that the cortical circuit did influence cross-orientation responses, but not in a manner consistent with cross-orientation suppression.

If cortex does not provide cross-column normalization, then one might ask if there is any evidence for recurrent influences within the cortical circuit to augment feed-forward input. In a second set of experiments, we looked for hallmarks of inhibition-stabilized activity, where providing additional drive to inhibitory neurons causes a “paradoxical” decrease in inhibitory responses instead of the expected increase due to the increased drive (Tsodyks et al., 1997; Ozeki et al., 2009; Sanzeni et al., 2020). These paradoxical responses presumably reflect a removal of recurrent excitation as inhibitory drive is increased. Using an inhibitory cell specific promoter (Dimidschstein et al., 2016), we provided optogenetic activation to cortical inhibitory neurons during visual stimulation. We observed a range of inhibitory neuron responses, including some that were “paradoxical” and others that were not.

## 3. Materials and Methods

### General design

All experimental procedures were approved by the Brandeis University Animal Care and Use Committee and performed in compliance with National Institutes of Health guidelines. Eleven adult ferrets (Mustela putorius furo, Marshall Farms; >P90, female) were used in total: five ferrets were used for patterned optogenetic experiments, four ferrets were used for optogenetic specificity experiments, and four ferrets were used for inhibitory optogenetic experiments. Females were used exclusively because co-housing male and female adult ferrets in the same space is stressful for the animals if they are not allowed to mate. For patterned optogenetic and optogenetic specificity experiments, AAV9.CamKIIa.hChR2(E123T/T159C).mCherry.WPRE.hGH was used to express ChR2 in neurons. Prior studies have found that high-titer AAV injections with the CamKIIa promotor causes expression that is biased towards excitatory neurons but expression is also found in inhibitory interneurons (Nathanson et al., 2009; Roy et al., 2016). AAV9-mDlx-ChR2-mCherry-Fishell-3, kindly provided by the Fitzpatrick lab at Max Planck Florida Institute, was used to express channelrhodopsin in interneurons (Dimidschstein et al., 2016).

### Virus injection

All virus injections were achieved by pre-treating ferrets with ketoprofen (1mg/kg, IM) and tramadol (2-5mg/kg, oral) on the morning of surgery. The ferrets were anesthetized with ketamine/xylazine cocktail (20-30mg/kg, 2- 3mg/kg) through IM injections and the anesthesia was maintained by additional injections of ketamine/xylazine (10-50% amount of ketamine/xylazine used during initial anesthesia). Atropine (0.16mg/kg) was given to reduce secretions. Ringer’s solution (2.75/ml/kg/hr) was given by subcutaneous injections to prevent dehydration. The body temperature was controlled and monitored by a thermostatic controller (TR-200, Fine Science Tools or PhysioSuite, Kent Scientific), and the EKG levels were continuously monitored.

Ferrets were held in a stereotaxic apparatus by two ear bars and a bite bar. The heads were shaved and sterilized by alternate applications of Betadine-soaked gauze and 70% isopropanol-soaked gauze three times. Bupivacaine (0.25- 0.5ml of 0.25% with a maximum does of 2mg/kg, IM) was injected around the incisions on the head. Head muscle and skin were retracted and a craniotomy, about 1-2 mm wide, was performed. A small durotomy was made with a 31-gauge needle on a cotton tip applicator. The glass pipettes used to inject viruses were pulled on a vertical puller (PC100, Narishige) and beveled to achieve a tip about 30um in diameter. Virus was delivered by a microinjection device (Nanoject, Drummond Scientific) with 22 pulses of 23 nl/pulse with 10 seconds intervals. To achieve a broad expression area (∼2.5 mm2), two or three locations were injected, two depths (300um and 500um below the brain surface) at each location, for each ferret. After the virus injection, the craniotomy site was covered with an Amniograft membrane (in some experiments) and the removed skull. The scalp incision was closed with non-absorbable sutures and the wound site was covered with Neosporin. Animals were returned to the cage with the rest of the litter after they were ambulatory. Analgesics and antibiotics were administered through 48 hours after surgery.

### Construction of ProjectorScope 2

The ProjectorScope 2 (Fig. 1, Fig. 1-1, 1-2) was built with several modifications of ProjectorScope 1 (Roy et al., 2016) to achieve wide patterned optogenetic stimulation and intrinsic signal imaging. Patterned light for optogenetic stimulation was generated by an LCD projector (NP3250W, NEC) and transmitted onto the brain surface by three single-lens reflex (SLR) lenses of the same type (Nikon, focal length 50 mm, f/1.2). The original projection lens was replaced with one of these SLR lenses to reduce misalignment between the projector and the rest of the optical system. Two juxtaposed lenses (L1; Thorlabs achromat, focal length 30mm, diameter 25mm) were used to minify the projection image to the appropriate size, ∼3mm by 3mm, in order to cover the exposed area in ferret V1. A dichroic mirror (DM; Semrock FF483/639) reflected light between 390-460 nm to activate ChR2, while allowing green and red light to pass to the CCD camera (Dalsa camera, 1M60) for intrinsic signal imaging. Intrinsic signal imaging was performed by providing 690-nm light (halide light, Lumen Dynamics, Xcite 200DC, with a filter, Semrock FF01-675/67-25) over the brain surface and taking images of the reflected light using the CCD, with an SLR lens (Nikon, focal length 135 mm, f/2.8) to bring the image into focus. The maximum power that the system can provide is approximately 10mW/mm2 measured at 475 nm by projecting a full-field, 100% contrast white image. ProjectorScope 2 allows three-dimensional adjustments for easier focus on a curved brain surface (details seen in Fig. 1-1, 1-2).

**Figure 1:**
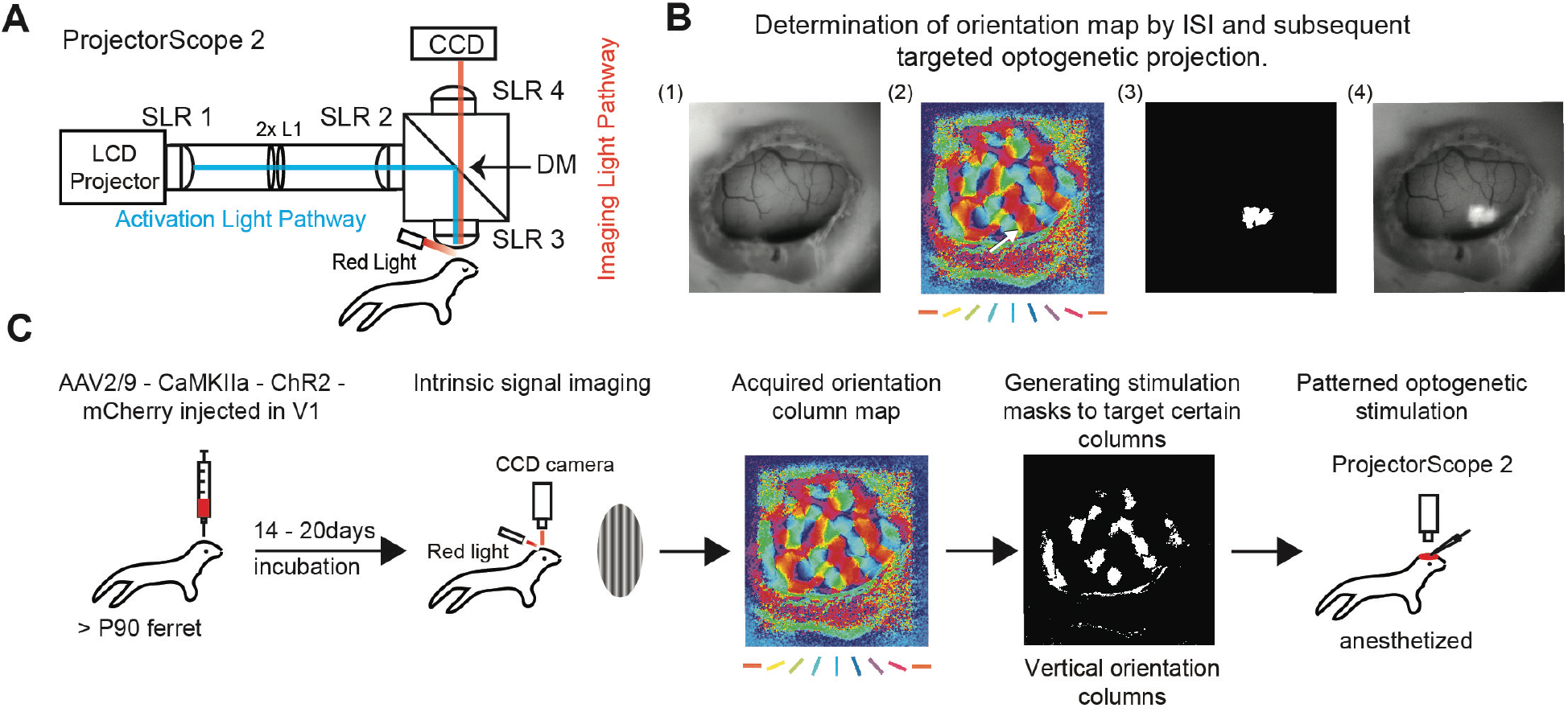
Optogenetic stimulation of functionally-identified cortical columns with ProjectorScope 2. **A**: ProjectorScope 2 construction. Patterned light for optogenetic stimulation is generated by an LCD projector and transmitted to the brain surface by three SLR lenses of the same type. Two juxtaposed lenses (2x L1) are used to minify the projection image to the appropriate size, ∼3mm by 3mm, in order to target several orientation columns in ferret V1. A dichroic mirror (DM) reflects light between 390-460 nm to activate ChR2, while allowing green and red light to pass to the CCD camera for intrinsic signal imaging (ISI). ISI is performed by providing brightfield red light over the brain surface and taking images of the reflected light using the CCD, with an SLR lens to bring the light into focus. **B**: Demonstration of intrinsic imaging and subsequent targeted optical stimulation. Determination of orientation map by ISI and subsequent targeted optogenetic projection. (1) Brain surface lit by room light. (2) The orientation column map acquired by ISI. The color key indicates the angle that a local region prefers. The arrow points to a region that will be used for targeted projection. (3) A projection mask based on the region pointed by the arrow in (2). (4) A raw image of the projection of the mask onto the brain surface. **C**: Experimental design. Viruses are injected to express ChR2 in V1 of adult ferrets, over 90 days old. After about three weeks, ISI is performed on the transfected ferrets to acquire an orientation column map. Then, the masks that target columns with certain orientation angles are generated based on the map.

**Figure 1-1.**
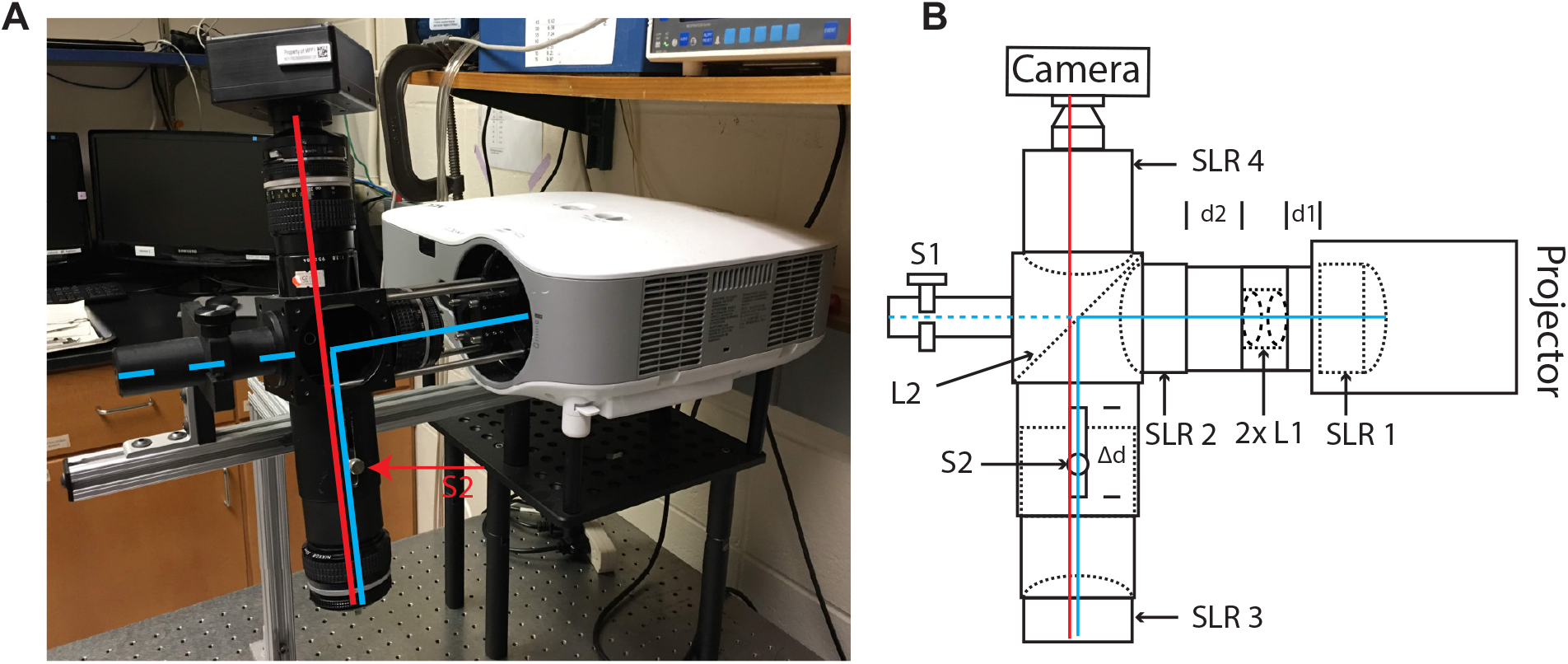
ProjectorScope 2. **A.** ProjectorScope 2 on the experimental table. The blue axis represents the direction of the outgoing light from the projector reflected by a dichroic mirror. The red axis represents the direction along which light travels from the brain surface to the camera during intrinsic signal imaging. The red arrow points to the tube that provides translation along the red axis using set screw S2. **B.** A schematic of ProjectorScope 2. SLR, single- reflex lens; S1, screw 1; S2, screw 2; 2x L1, two closely placed achromat lenses; L2, dichroic mirror; d1=30mm; d2=45mm; Δd = 6cm.

**Figure 1-2.**
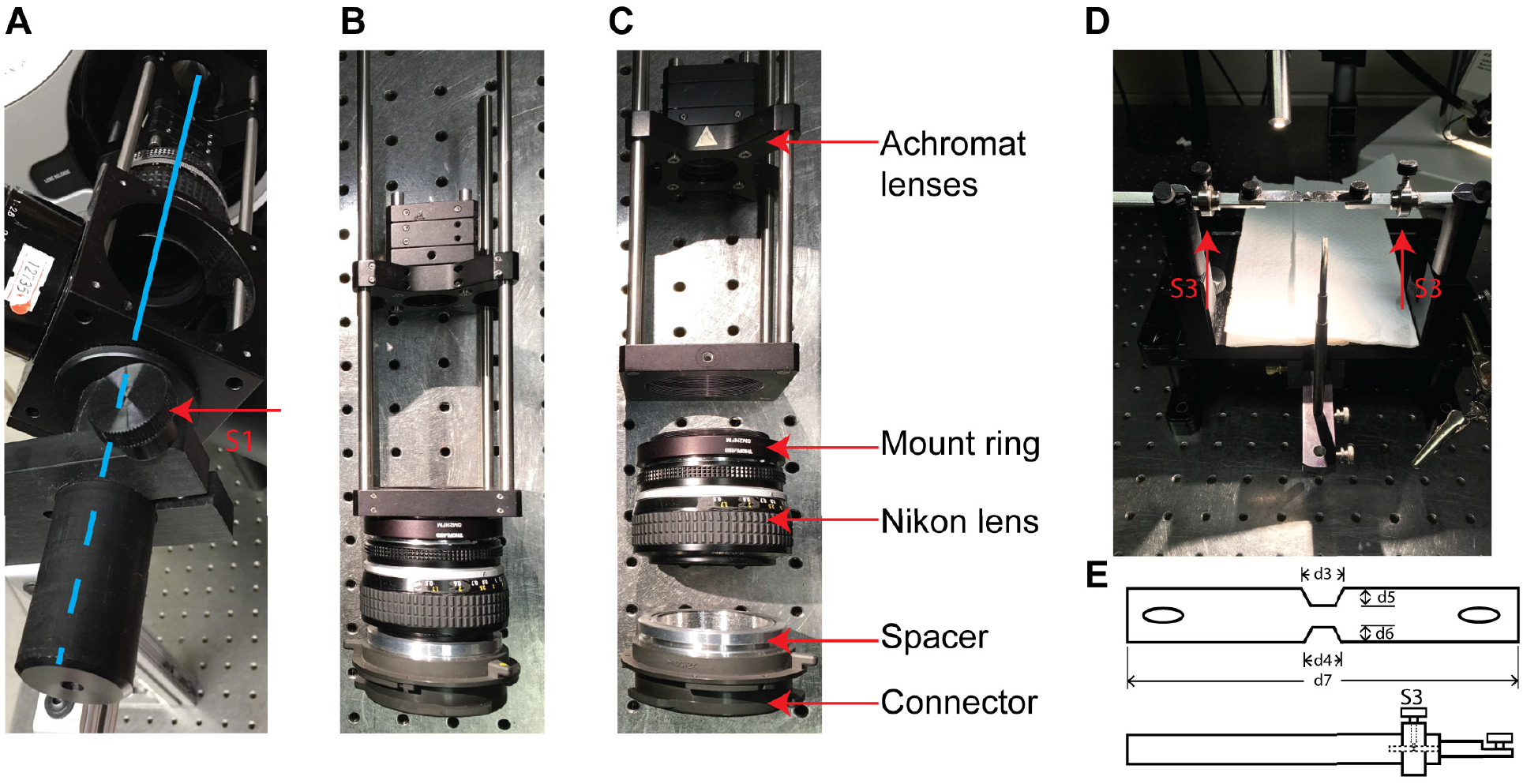
Components of ProjectorScope 2 and the custom head-plate and head-bars. **A.** The cylindrical bar that allows the cube to rotate around the blue axis. The knob S1 controls the rotation of the bar. **B.** The replaced projection lens with all components connected. **C.** A disconnected version of B, which separately displays the connector to the projector, the spacer, the Nikon lens, and the custom-made mount ring that connects the Nikon lens to the Thorlabs rectangular connector. **D.** The rotatable headplate. The red arrows point to the rotatable parts, S3, that can adjust the animal’s head pitch angle. Combining the controllable parts in S1, S2, and S3, ProjectorScope 2 allows three-dimensional adjustments for easier focus on a curved brain surface. **E.** Schematics of the head-plate and the head-bar. The head-plate, 1mm thick, has two half-hexagon notches to easily fit the curved skull surface. d3=7mm, d5=2mm, d4=5mm, d6=1.5mm, d7=6.5cm. The head-bar provides rotation by adjusting S3.

### Non-survival surgery

About 4 weeks after the virus injection, the ferrets were sedated with ketamine (20mg/kg, IM). Atropine (0.16mg/kg, IM) and dexamethasone (0.5mg/kg, IM) were administered to reduce bronchial and salivary secretion and to reduce inflammation, respectively. The animal was anesthetized with a mixture of isoflurane, oxygen, and nitrous oxide through a mask while a tracheostomy was performed. Animals were then ventilated with 1.5%–3% isoflurane in a 2:1 mixture of nitrous oxide and oxygen. A cannula was inserted into the intraperitoneal cavity for delivery of neuromuscular blockers and Ringer’s solution (3 ml/kg/hr), and the animal was placed in a custom stereotaxic frame that did not obstruct vision. The head was fixed with a custom head plate that allowed pitch adjustments for imaging. All wound margins were infused with bupivacaine. Silicone oil was placed on the eyes to prevent corneal drying. A craniotomy (4 by 4mm) was made in the right hemisphere centered around the virus injection site, and the dura was removed with a 31-gauge needle. A few drops of liquid agarose were applied on the exposed brain surface, and, while the agarose was still liquid, a pre-drilled coverslip (a hole of about 700um in diameter was drilled through the coverslip) was mounted on top of the craniotomy area and held until the agarose became solid (Levy et al., 2012). The coverslip edge was secured using cyanoacrylate glue and excess agarose on the coverslip was removed. Next, the ferrets were paralyzed with the neuromuscular blocker gallamine triethiodide (10–30 mg/kg/hr) through the intraperitoneal cannula to suppress spontaneous eye movements, and the nitrous oxide-oxygen mixture was adjusted to 1:1. The animal’s ECG was continuously monitored to ensure adequate anesthesia, and the percentage of isoflurane was increased if the ECG indicated any distress. Body temperature was maintained at 37°C.

### Visual stimulation

Visual stimuli were created in MATLAB with the Psychophysics Toolbox on a Macintosh Pro running OSX and displayed on a Sony monitor (GDM-520). The monitor was placed 35cm in front of the ferret. Stimuli were full field sine wave gratings with 0.15 spatial frequency and 4Hz temporal frequency.

For intrinsic signal imaging experiments, 100% contrast, bidirectional grating stimuli with orientations varied from 0° to 135° with a step of 45° were played. Each orientation condition was repeated 20 times with 10s interstimulus-interval and 5s stimulus duration.

For the optogenetic experiments, visual stimuli were 100% contrast, full-field, with 0.15 spatial frequency, 4Hz temporal frequency with 8 cycles and repeated 5 times with 3-5s inter-stimulus-interval. To measure the orientation selectivity, the orientations were varied from 0° to 157.5° with a step of 22.5°. To measure responses to visual contrast, the orientation was fixed at the preferred angle and the contrasts were varied as 16%, 32%, 64%, or 100%.

Intrinsic signal imaging. Intrinsic signal imaging was performed for some optogenetic experiments to obtain the orientation column maps. With ProjectorScope 2, 690-nm light illuminated the brain surface and the reflected light from the brain surface was captured by the camera. The images were acquired at 30 Hz with custom software in LabVIEW and a National Instruments PCI-1426 acquisition board. The raw images were averaged for every 0.5s and the image that was 0.5s before the onset of the stimulus was used as the baseline image. The single condition images representing responses for individual orientations were averaged over all repetitions and the orientation column map was generated by calculating the vector summation of responses in the single condition images:

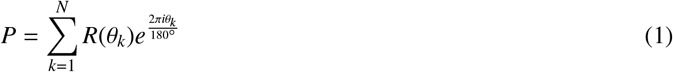

where *R*(θ*_k_*) is the responses in a single condition image for a certain orientation stimulus, and P represents the response of each pixel in the map as a vector summation in the complex plane.

### Electrophysiology

In some optogenetic experiments and optogenetic specificity experiments, single barrel carbon fiber electrodes were used (E1011, Kation Scientific). Such carbon fiber electrodes have very small tips of about 5um in diameter so they minimize damage to the brain tissue and do not cast shadows over the stimulated brain area. One electrode was inserted through the hole on the coverslip into ferret V1, and was lowered to a depth that ranged from 100um to 400um below the brain surface. The signals were amplified by RHD2132 and collected by the RHD2000 evaluation board (Intan Technologies).

In the inhibitory optogenetic experiments, a custom 32-channel probe (S-probe, Plexon) was used. Instead of using pre-drilled coverslips, the probe was positioned to just touch the brain surface and 2%-3% agarose was applied to prevent brain pulsation. After the agarose stabilized the brain, the probe was then lowered until all pads were inserted (1000-1200um in the diagonal direction). Mineral oil was applied to the agarose at regular intervals to prevent the agarose from drying.

### Optogenetic receptive zone

We characterized the optogenetic receptive zone (ORZ) for each patterned optogenetic experiment. To measure the spatial range of the effective optogenetic stimulation, we projected small dots, ∼750um in diameter, in a randomized fashion tiling across the entire projection area, ∼3 by 3 mm2, onto the ferret primary visual cortex that had ChR2 expression. We determined whether a cell’s response at any of the stimulus positions was significantly different from the response to a “blank” stimulus by performing an ANOVA test (P<0.05). Responses were fitted by a bivariate Gaussian function to estimate the region over which a cell was strongly activated :

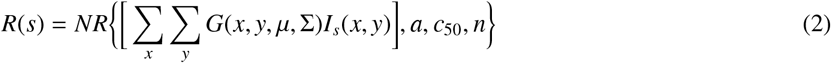

Where *I_s_*(*x*, *y*) is the intensity at point x,y for stimulus s, *G*(*x*, *y*, µ, Σ) is the bivariate Gaussian with mean µ and covariance matrix Σ, and *NR*(*c*, *a*, *c*_50_, *n*) is the Naka-Rushton function:

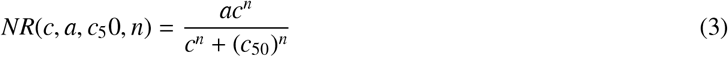

Where a is the maximum cell response, c is the stimulus intensity, and *c*_50_ is the intensity of a stimulus that produces half of the maximum response. Variables a, *c*_50_, n, µ, Σ were used as free parameters for the fit. The size of the ORZ was taken to be the full width at half-height (FWHH) along the major and minor axes of *G*(*x*, *y*, µ, Σ).

### Patterned optogenetic stimulation

In patterned optogenetic specificity experiments, stimulation masks targeting specific orientation columns were generated based on the map acquired during intrinsic signal imaging (custom software, MATLAB). The preferred orientation of the recorded neuron was identified as the orientation that evoked the strongest visual responses from the orientation tuning curve. Visual contrast tuning curves were initially measured using 16%, 32%, 64%, or 100% contrast. The optogenetic light intensity tuning curve were measured by projecting the mask of the ORZ with varying image intensities, 20%, 40%, 60%, 80%. The measurements of these tuning curves were repeated by changing contrasts or intensities until comparable levels between visual and optogenetic responses were found, which was usually achieved after 2-3 iterations. Five levels of visual contrasts or light intensities, including 0% contrast and 0% intensity, were chosen for each cell. For the combined visual and patterned optogenetic stimulation, each visual stimulus was paired with an optogenetic mask stimulus and each pair started at the same time with the optogenetic stimulation lasting for 1s and visual stimulation lasting for 2s. The orientation column masks for a given angle were made by including all pixels in the intrinsic signal imaging map that matched the specified angle within some tolerance. The tolerance (or thickness) of the column masks was varied by changing the range of orientation angles that each column mask contained: 15°, 30°, or 45° tolerance. We found that a tolerance of 15° was too small for some cells to evoke enough optogenetic responses and that a tolerance of 45° was too big to create distinctly complementary patterns of inputs, so, in the analysis here, we only report the results based on 30° masks. The stimulation order was randomized.

For all-optogenetic stimulation (Fig. 3A), the masks were created to target either the preferred orientation columns that were revealed in the orientation columns map, the orthogonal orientation columns, or both. For hybrid stimulation (Fig. 3B), the masks were created to target the orthogonal orientation columns and the visual stimulus presented the preferred orientation. For the optogenetic specificity experiments, only the orientation columns inside the ORZ were included and the masks were created to target the orientations from 0° to 157.5° with a step of 22.5° and a tolerance of 15°.

### Pharmacological blocking experiments

To test the hypotheses of optogenetic specificity, an NMDA antagonist (DL-2-Amino-5-phosphonopentanoic acid, APV, 1mM- 5mM) and an AMPA antagonist (2,3-Dioxo-6-nitro-1,2,3,4-tetrahydrobenzo[f]quinoxaline-7-sulfonamide, NBQX, 100uM-1mM) were used to block NMDA and AMPA receptors. For these experiments, the coverslips were pre-drilled with two holes that were 2-3mm apart, one for the carbon fiber electrode and one for the blocker pipette. A pulled glass pipette similar to the dimension used in virus injections was used to apply the blockers and the blocker solution was delivered by Nanoject, with 69nl/pulse, 4 pulses/min, for 14 min, for a total volume of ∼3.8ul. The effectiveness of the blockers was tested by examining visual responses. Neurons were said to give significant responses if any response exceeded 3 standard errors above the background response. Light intensities of 80 and 100% were averaged together to compute tuning curves.

### Data analysis

Extracellullar data was extracted using 4-5 standard deviations as the threshold and then clustered using K-means or KlustaKwik based on either two-point features (for the single barrel carbon fiber electrode recordings) or the principal components (for the S-probe recordings). In the patterned optogenetic experiments, the firing rates were calculated as spike counts from the 0.5s-1s after the onset of the stimuli divided by 0.5s duration, because the firing rates reached steady state after 500ms of stimulation. In the inhibitory optogenetic experiments, the calculations of firing rates were based on the entire 1s duration of paired stimulation.

To analyze the optogenetic specificity, the specificity index is defined as 1-circular variance (Ringach et al., 2002; Mazurek et al., 2014), calculated as:

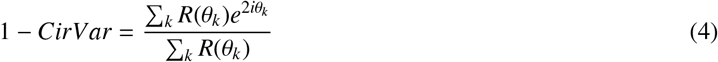

Where *R*(θ*_k_*) is the response to angle θ*_k_*. The comparison between the specificity before and after blockers applications was based on two-sample t-test.

To summarize the effects of visual and optogenetic stimulation on recorded neural responses, we fit a divisive normalization model (Carandini and Heeger, 2011) to the responses of each cell. The model describes the firing rate response *R* as:

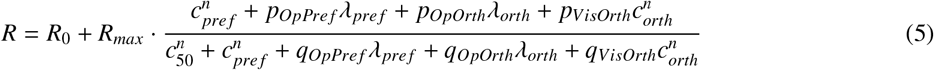

where *R*_0_ is the baseline firing rate, *R_max_*is the maximum firing rate above baseline, and the remaining parameters describe how different stimulus inputs drive and normalize the response. *c_pre f_* is the contrast of the visual stimulus at the cell’s preferred orientation, *c_orth_* is the contrast of the visual stimulus at the orthogonal orientation, λ*_pre f_* is the intensity of optogenetic stimulation of preferred-orientation columns (normalized to 1 at maximum laser power), and λ*_orth_* is the intensity of optogenetic stimulation of orthogonal-orientation columns. The exponent *n* controls the steepness of contrast responses and *c*_50_ is the semi-saturation contrast. Note that the preferred visual drive *c^n^* appears in both numerator and denominator without separate *p* and *q* coefficients; the *p* and *q* parameters are only needed for inputs beyond the preferred visual drive, because the preferred visual response is the reference against which all other inputs are compared.

The parameters *p_OpPre f_*, *p_OpOrth_*, and *p_VisOrth_* determine the degree to which optogenetic stimulation of preferred columns, optogenetic stimulation of orthogonal columns, and visual stimulation of the orthogonal orientation, respectively, activate the recorded cell (numerator contributions). The corresponding parameters *q_OpPre f_*, *q_OpOrth_*, and *q_VisOrth_* determine the degree to which these same inputs suppress the recorded cell through divisive normalization (denominator contributions). The parameters *c*_50_ and *n* were shared across all stimulus conditions for a given cell, while *R*_0_ and *R_max_* were fit per cell. All parameters were constrained to be non-negative.

The ratio of each activation parameter to its corresponding suppression parameter has a natural interpretation as an asymptotic ceiling.

The fits were done in two stages, first considering the visual-only parameters (25 measurements), followed by fits for the *p* and *q* variables using the optogenetic observations (visual-preferred with optogenetic-orthogonal, and optogenetic-only) with 48 measurements. In all there were ten free parameters per cell for 73 mean response measurements over the various conditions. Responses were fit in their measured units (spikes s^−1^), without blank subtraction or per-cell rescaling, with the optogenetic drives λ*_pre f_* and λ*_orth_* normalized to the maximum laser power across the data set so that each lay in [0, 1]. Beyond the non-negativity constraint, parameters were bounded as follows: *c*_50_ ∈ [10^−4^, 1], spanning the full range of normalized contrast, so that the upper bound corresponds to semi-saturation at the maximum contrast presented; *n* ∈ [0.5, 4]; the activation and suppression weights *p_OpPre f_*, *p_OpOrth_*, *p_VisOrth_*, *q_OpPre f_*, *q_OpOrth_*, *q_VisOrth_* ∈ [0, 10]; and *R*_0_, *R_max_* ∈ [0, 200] spikes s^−1^.

Parameters were estimated by minimizing the sum of squared residuals between the model prediction and the measured mean responses, using constrained nonlinear optimization (interior-point algorithm; up to 1000 iterations and 5000 function evaluations per run). To reduce sensitivity to local minima, each cell was fit with a multi-start procedure comprising 50 independent optimizations. The first optimization began from a fixed initialization (*c*_50_ = 0.2, *n* = 2, all *p* and *q* weights = 1, with *R*_0_ and *R_max_* initialized from the minimum and the range of that cell’s responses, respectively); the remaining 49 optimizations began from initial parameter vectors drawn independently from a uniform distribution over the bounded parameter space. The parameter set with the lowest residual sum of squares across all starts was retained as the fit for that cell, and the random number generator was seeded for reproducibility. Goodness of fit was quantified as the coefficient of determination (*R*^2^), computed both overall and separately for each stimulus condition.

To determine whether optogenetic stimulation contributed significantly to each cell’s responses, and whether optogenetic stimulation of orthogonal-orientation columns contributed beyond that of preferred-orientation columns, we compared the full ten-parameter model against reduced models nested within it using an extra-sum-of-squares (*F*) test. Each reduced model was obtained by constraining a subset of the optogenetic weights to zero and was then refit to that cell’s responses with the identical multi-start procedure, so that its residual sum of squares reflected the true optimum of the constrained model rather than the full fit with parameters clamped. To test for any optogenetic contribution, we fit a reduced model in which all four optogenetic weights *p_OpPre f_*, *p_OpOrth_*, *q_OpPre f_*, and *q_OpOrth_* were fixed at zero (six free parameters); to test specifically for a contribution of orthogonal-column optogenetic stimulation, we fit a second reduced model in which only *p_OpOrth_* and *q_OpOrth_*were fixed at zero (eight free parameters), leaving the preferred-column optogenetic weights free. For each comparison we computed

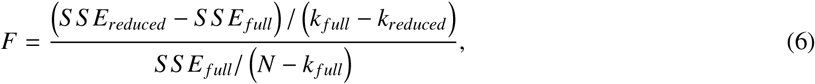

where *S S E _f ull_* and *S S E_reduced_* are the residual sums of squares of the full and reduced fits, *k _f ull_* = 10 and *k_reduced_* is the number of free parameters in the reduced model, and *N* = 73 is the number of mean response measurements per cell. Under the null hypothesis that the constrained weights are jointly zero, *F* follows an *F* distribution with *k _f ull_* − *k_reduced_*, *N* − *k _f ull_* degrees of freedom, giving *F*_(4,63)_ for the test of any optogenetic contribution and *F*_(2,63)_ for the test of an orthogonal-column contribution; cells with *p* < 0.05 were taken to show a significant effect. Cells that showed no significant optogenetic influence in the 4 parameters were excluded, and cells that showed significant preferred optogenetic stimulation but did not show significant orthogonal optogenetic activation were noted in the figures.

### Ring model simulations

The ring models consisted of 180 E and I cells, placed along a ring. Each position was given a label θ*_i_*= 0. . . 179°.

The steady-state response of each cell was given by the equation from Rubin et al. 2015:

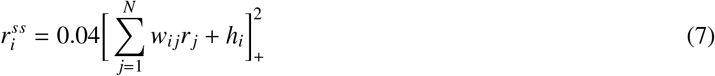

where *w_i j_* is the connection from neuron *j* to neuron *i*, and *h_i_* is the sum of visual (*h_visual_*) and optogenetic (*h_opto_*) input to neuron *i*. The connections from e-to-e neurons had a weight *W_EE_* that fell off in a Gaussian manner with angular distance around the ring as σ*_EE_*. Similarly, connections from i-to-e had a value *W_EI_* that fell off with σ*_EI_*, connections from e-to-i had a value *W_IE_* that fell off with σ*_IE_*, and i-to-i connections had a weight *W_II_* that fell off with σ*_II_*. The following differential equation was solved numerically using Euler’s method:

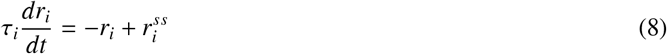

where τ*_i_* is 200ms for excitatory cells and 100ms for inhibitory cells. Visual stimulation at an angle θ was delivered to each neuron with a strength of 1 and a Gaussian falloff of 30°.

The details of the connections and optogenetic input differed by model. For Model 1, *W_EE_* was 0.018, *W_EI_* was -0.0094, *W_IE_* was 0.0171, *W_II_* was -0.0073, and σ*_EE_* = σ*_IE_* = σ*_EI_* = σ*_II_* = 32°. Optogenetic input was delivered to each neuron with a strength of 1 and a Gaussian falloff of 45° from the angle of columnar stimulation θ.

For Model 2, the parameters were the same as Model 1, except that the Gaussian portion of the optogenetic inputs were decreased (multiplied by 0.7) and a constant, angle-independent input across all excitatory and inhibitory neurons (weight 0.28) was added. σ*_EE_* = σ*_IE_* = σ*_II_* = 10°, while σ*_EI_* was dropped to 15°. Normalization was provided on the visual input side (to both excitatory and inhibitory neurons) with the equations:

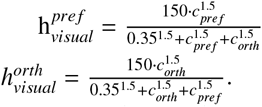

For Model 2L, the parameters were the same as Model 2, except that the visual input was returned to the linear input of Model 1. The purpose of Model 2L is simply to visualize the within-cortex actions of Model 2.

### Heterogeneous network simulations

To simulate the impact of optogenetic stimulation in a heterogeneous network with variable optogenetic activation and variable synaptic weights, we simulated a column with 2 populations with *N* neurons in it, each with steady-state rate ⃗*r* given by:

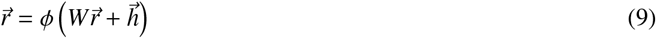

Where the function ϕ is a modified rectified power law function ϕ(*x*) = (α*x*)*^n^*/(1 + (*x* + α)*^n^*), which saturates for *x* → ∞ and behaves similarly to the standard rectified power law or small values of x. We chose alpha =35 and *n* = 2 The connectivity elements are sparse with sparsity *p* = 0.3. The nonzero elements of W are *w_ee_*=1, *w_ei_*=1.1, *w_ie_*=0.89, *w_ii_*=0.91 and they scale as 1/N with the network size.

The external inputs to each neuron *h*^α^ have a baseline component and an optogenetic input, such that

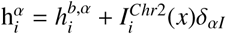

The baseline was uniformly distributed *h*, centered in µ*_h_b* = 10 and with a width σ*_h_b* = 6 for both cell types.

The optogenetic input only affects inhibitory neurons and is defined below.

### Model of Optogenetic Perturbations

There were two main sources of photocurrent heterogeneity, one being light dispersion through the tissue and the second one being the number or ChR2 channels that are expressed in each cell. Regarding the first one the amount of light that reaches the cells decays exponentially with distance (Yona et. al. 2016). We assumed that the recorded neurons are homogeneously distributed in the z axis of the probe. The light that arrived to each neuron was given by

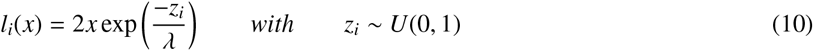

Where *x* is the light intensity at the surface, *z_i_*is the distance of the cell *i* from the surface, and λ is the spatial scale (We took λ = 2). We assumed that, each channel had an input output function that is given by a Hill Equation and that each channel had a different threshold, given by *l*^0^

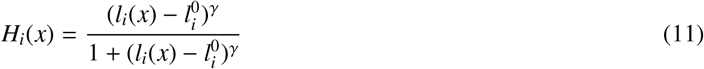

With *l*^0^ = *l_i_*(*x*_0_), with *x*_0_ ∼ N(µ*_x_*_0_, σ*_x_*_0_). The values were chosen to be µ*_x_*_0_ = −0.05 while σ*_x_*_0_ = 0.001. The second source of heterogeneity is the number of ChR2 channels that are expressed with the virus, that we assume to be gaussian distributed. In the simulations, we didn’t specify the number of channels, but we used a normalized number distributed as κ*c* ∼ N(µ*_c_*, σ*_c_*) with µ*_c_* = 72 and σ*_c_* = 8

Each individual channel contributed independently and identically to the photocurrent such that the total input to the cell was

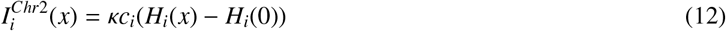

Where the substraction by *H_i_*(0) guaranteed that the effect of the laser at zero intensity (x=0) was 0.

## 4. Results

### Direct stimulation to test normalization mechanisms in cortex

Visual cortical neurons exhibit sublinear responses to the simultaneous presentation of two grating stimuli (Adelson and Movshon, 1982; Morrone et al., 1982; Morrone et al., 1987; DeAngelis et al., 1992; Heeger, 1992; Carandini et al., 1997; Popovic et al., 2018), Because at least some of this suppression is present in the inputs to the cortex that arise from LGN (Freeman et al., 2002; Li et al., 2006; Priebe and Ferster, 2006; Barbera 2022), we designed an experiment to directly provide input to different regions of the cortical circuit itself, in order to separate cortical contributions to normalization from those of its inputs.

To do this, we updated a custom-made optical system (Roy et al., 2016), now called ProjectorScope 2 (Fig. 1A), that allowed us to use intrinsic signal imaging (Blasdel and Salama, 1986; Grinvald et al., 1986) to identify the locations of orientation columns and also allowed us to provide patterned optogenetic stimulation to the cortical surface in a manner that matched orientation columns or any other desired pattern (Fig. 1B). In our first set of experiments, we injected viruses to express a variant of ChR2 (Boyden et al., 2005; Berndt et al., 2011) in neurons and prepared the ferrets for intrinsic signal imaging. After the orientation column map was acquired, optogenetic stimulation masks that targeted certain orientation columns were calculated based on the empirical map, and masked images were subsequently projected onto the V1 surface to stimulate the corresponding columns (Fig. 1C).

### Single-column optogenetic stimulation causes wide spread of activity due to horizontal connections

A key requirement of our experiment was to demonstrate that we could provide distinct input to different groups of nearby cortical neurons. While it was clear from visual inspection of the camera image that the light stimulus was illuminating distinct portions of the cortical circuit, several outcomes were possible at the neural level. First, providing direct input to a small column of neurons might only activate the neurons that were stimulated. Second, our direct optogenetic input might be restricted to distinct groups of neurons, but that activity could spread across columns through cortical synaptic connections; in this case, we would need to perform additional experiments with synaptic blockers to show that optogenetic inputs were being provided to distinct groups of neurons. Third, because axonal projections extend for millimeters and dendritic trees extend for a few hundred microns across the cortical surface, activation of these axons and dendrites might be sufficient to drive spiking in cell bodies over a wide area, which would mean that we would be unable to provide inputs to distinct groups of nearby neurons even with precise optical stimulation of the cortical surface.

Using electrodes, we characterized the optogenetic receptive zone (ORZ) of single neurons by stimulating the brain surface using circular dots (≈ 750µ*m* in diameter) in a randomized fashion. A 2-D Gaussian fit was performed on the recorded optogenetic responses over the cortical surface to delineate the local region that could be effectively activated by light (Fig. 2A). The ORZ could therefore be described as an elliptical shape comprising the interior 63%-tile of the fit (Roy et al., 2016).

**Figure 2:**
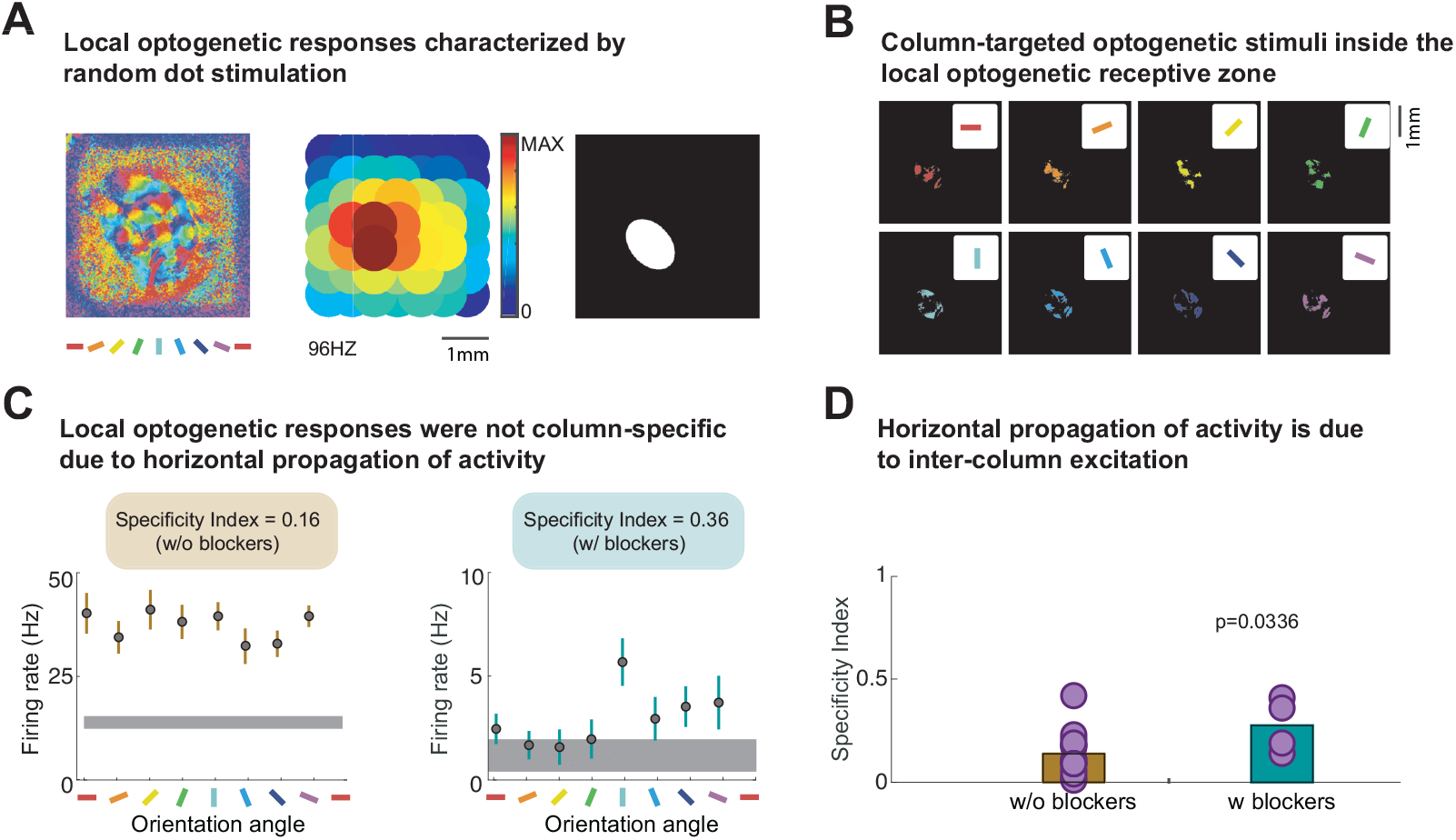
Although optogenetic activation of cortex produced local activity, this activity did not respect boundaries of orientation columns, due to horizontal propagation of activity. **A**: The orientation column map (left); the responses of a single unit to optogenetically-provided spot stimuli (750 µm in diameter) with the heat map indicating the response intensities with scale max of 96Hz (middle); the elliptical optogenetic receptive zone (ORZ) characterized by two-dimensional Gaussian fit of the responses to spot stimulation (right). **B**: The masks used to test the orientation column specificity of the optogenetic stimulation were generated inside the optogenetic receptive zone and targeted orientation columns of varying angles with steps in 22.5 degrees. **C**: Left: A neuron’s responses to the orientation specificity test before synaptic blockers. Right: A different neuron’s response to the orientation specificity test after synaptic blockers. The orientation angles shown on the x-axis are the angles of the corresponding column masks. **D**: The average specificity index of neurons that exhibited responses > 3 standard deviations above the blank (n=14 without blockers; n=4 with blockers) after synaptic blockers are applied is higher than that when no blockers are used (P<0.05). These results imply that ChR2 stimulation of the cortex in animals of this age activates several adjacent orientation columns, in part due to synaptic propagation of signals across the cortex.

We found that ORZs were larger in these adult ferrets (about the size of a ferret hypercolumn) than in our previous work in young ferrets (Roy et al., 2016), so we created stimuli to test the specificity of activation of different orientation columns that were restricted to the ORZ (Fig. 2B). In this manner, we examined the “tuning” of each neuron to optogenetic activation of various orientation columns within the ORZ. We found that responses were quite non-specific, indicating that cortical activity spread substantially across orientation columns (Fig. 2C, left). There are two possible sources for this spread: either our optical stimulus itself caused widespread activation, or direct optogenetic activation was restricted to specific regions of the cortical surface and the spread of activity was due to activity within the cortical network.

To differentiate the two scenarios, we applied synaptic blockers (NBQX and DL-AP5 to block AMPA and NMDA receptors, respectively) and measured the specificity index (1- circular variance) before and after the application of synaptic blockers. We found that neurons exhibited more specific responses to the preferred-column activation in the presence of synaptic blockers (Fig. 2D, t-test, P = 0.033). This suggests that some of the non-specific column-based optogenetic stimulation is due to the spread of evoked neural activity via connections intrinsic to the visual cortex, consistent with previous studies in tree shrew (Huang et al., 2014). Nevertheless, the fact that responses evoked with optogenetic stimulation in the presence of synaptic blockers were more local showed that we were able to provide somewhat selective input to distinct regions.

### Visual, mixed visual and optogenetic stimulation, and paired optogenetic stimulation have different normalization properties

Having established that we could provide distinct independent inputs to different groups of cortical neurons, we examined the cortical contributions to normalization by comparing how the response to the simultaneous presentation of a pair of stimuli was related to a simple linear sum of the responses to the stimuli independently. In all, we compared cortical integration under three conditions: visual, combined visual and optogenetic, and all optogenetic (Fig. 3).

**Figure 3:**
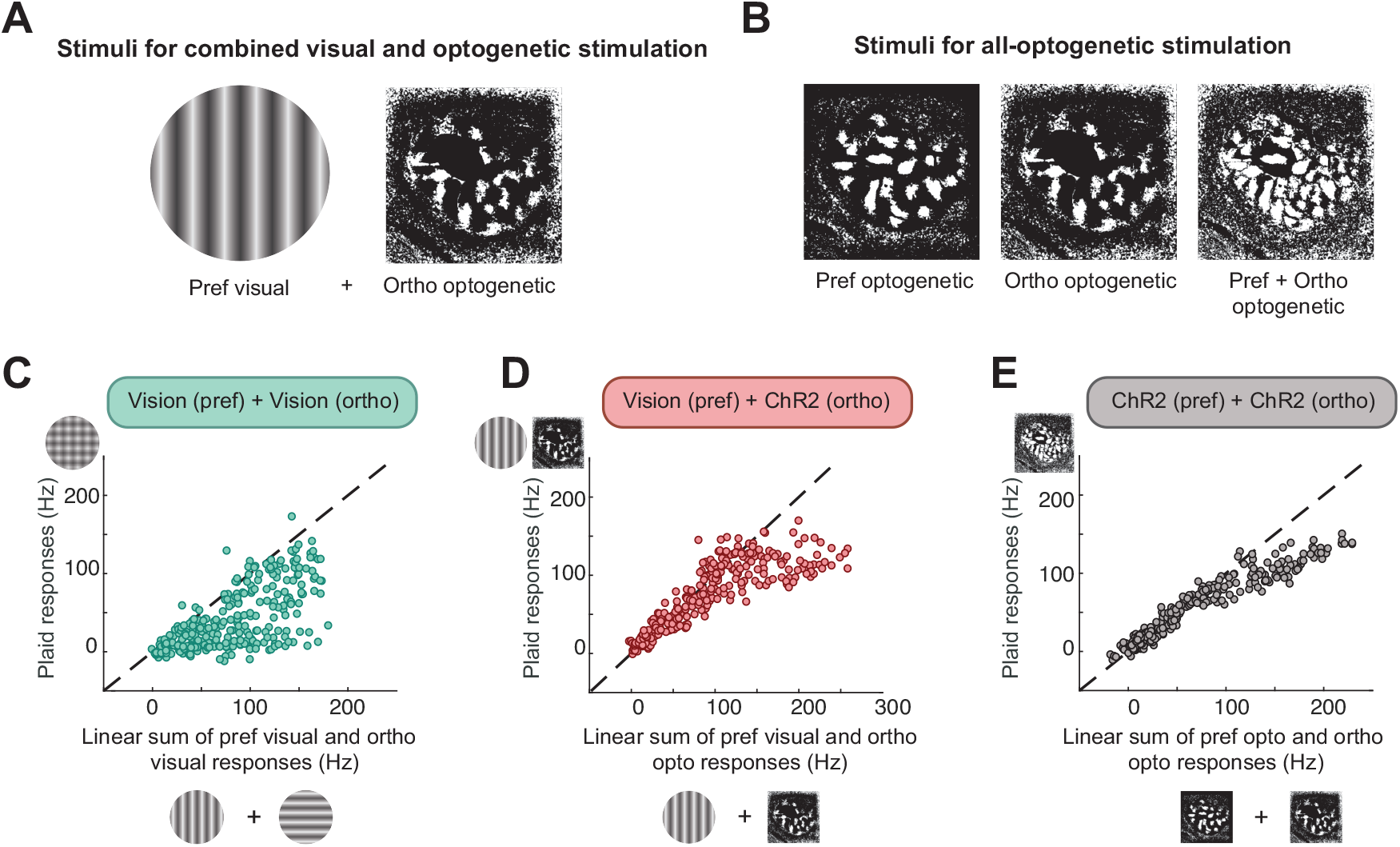
Neurons exhibit sublinear responses to combined visual and optogenetic stimulation. **A**: The stimuli used for hybrid stimulation. The visual stimulus provides the preferred orientation of the recorded neuron and the projection mask provides optogenetic modulation. **B**: Optogenetic stimuli for activation of neuron recorded with an electrode. Left) A strong optogenetic stimulus that includes the electrode region and all columns that prefer the same orientation (within 30°) as the recording site. Middle) A modulating stimulus that excludes the 1-sigma optogenetic receptive zone but includes all columns that prefer the orthogonal orientation (within 30°) as the recording site. Right) Combined stimuli. **C**: Single-unit responses to paired visual stimulation of preferred and orthogonal orientations, compared to the linear sum of preferred and orthogonal responses when those stimuli were presented alone. For each cell (N=23 cells, from 5 animals), responses to 16 combinations of preferred and orthogonal orientations are shown (4 contrast levels each). Responses are clearly sublinear for stimuli that evoked low or high firing rates. Dashed line is unity line. **D**: Single unit responses to paired visual and optogenetic stimulation. For each cell, responses to 16 combinations of visual preferred orientation (4 contrast levels) and optogenetic stimulation of the orthogonal orientation columns (4 drive levels) are shown. For stimuli that evoked low firing rates, paired stimulation exhibited mostly linear summation, but this summation became more sublinear for stimuli that evoked higher firing rates. **E**: Single unit responses to optogenetic stimulation of a cell’s preferred columns and orthogonal columns. For each cell, responses to 16 combinations of optogenetic drive (4 drive levels for preferred stimulation and orthogonal stimulation) are shown. For stimuli that evoked low firing rates, paired stimulation produced a response that was nearly the same as the linear sum of the two component stimuli when presented alone. For stimuli that evoked larger responses, summation became non-linear.

**Figure 3-1.**
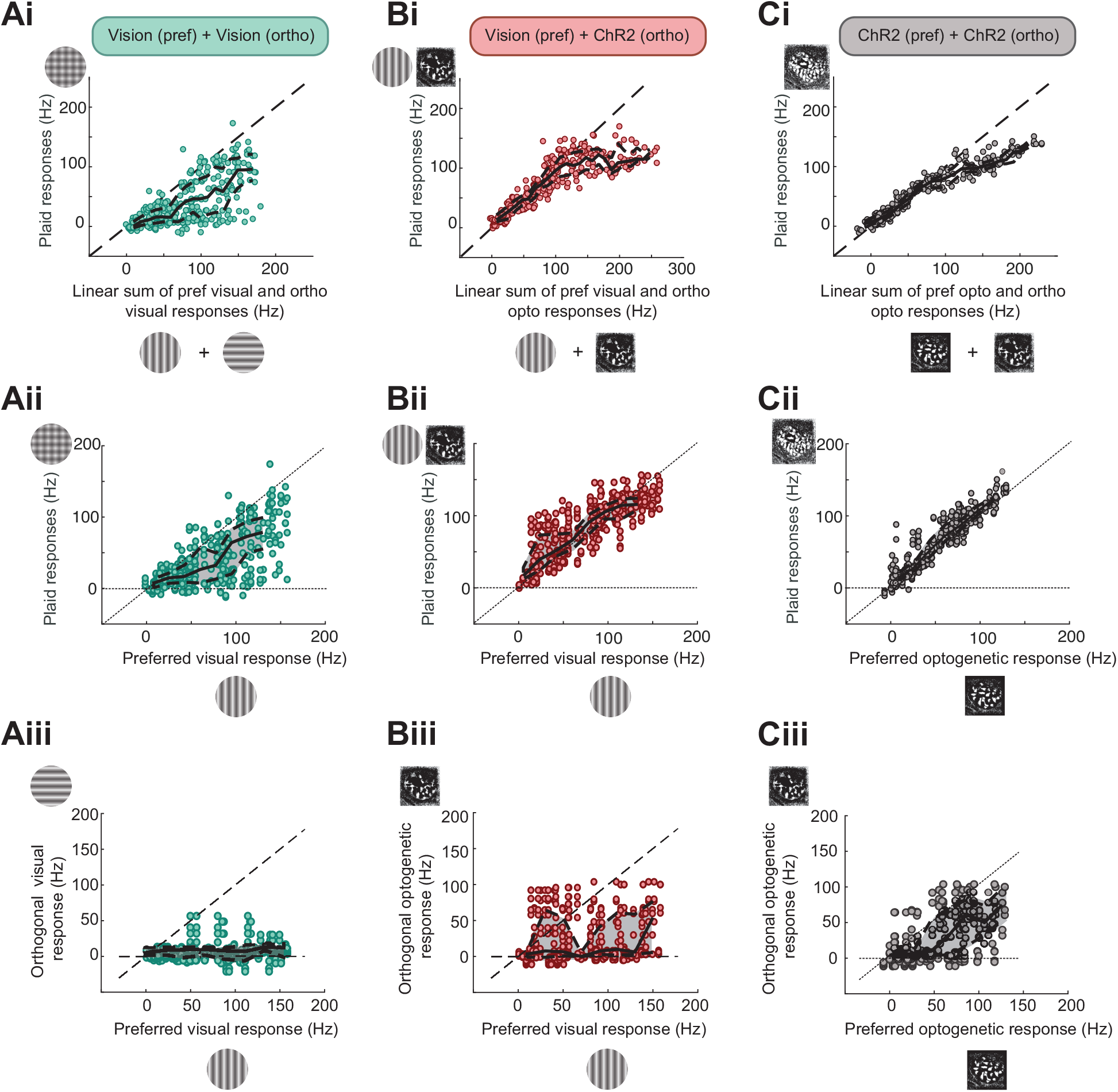
Neurons exhibit sublinear responses to combined visual and optogenetic stimulation. A) Single-unit responses to visual stimulation. i) Response to paired visual stimulation of preferred and orthogonal orientations, compared to the linear sum of preferred and orthogonal responses when those stimuli were presented alone. For each cell (N=23 cells, from 5 animals), responses to 16 combinations of preferred and orthogonal orientations are shown (4 contrast levels each). Responses are clearly sublinear for stimuli that evoked low or high firing rates. Solid black line indicates sliding median, dashed lines indicate 25-75%-tile range. ii) Paired preferred and orthogonal stimulation responses plotted against the response to the preferred orientation. iii) Orthogonal orientation responses plotted against preferred responses. Orthogonal visual stimuli generally produce poor responses. B) Same, except the preferred stimulus is delivered visually and the orthogonal stimulus is delivered optogenetically to the cortical surface (see Figure 3). When orthogonal optogenetic stimulation is paired with visual stimuli, the magnitude of the orthogonal stimulation does not depend on the magnitude of the response to visual stimulation (iii), and interactions are linear at low response intensities and become non-linear at higher response intensities (i, ii). C) Same, except preferred stimulus and orthogonal stimulus is delivered optogenetically (see Figure 3). Optogenetic preferred stimulation responses and optogenetic orthogonal responses are correlated (iii), presumably because optogenetic expression varies from animal-to-animal.

In the first condition (all visual), we examined the classic phenomenon of cross-orientation suppression by comparing the sum of the responses of neurons to visual stimulation at the preferred orientation or at the orthogonal orientation to the response to a visual “plaid” of the two orientations presented together, at a variety of stimulus contrasts. We compared these results to stimulation with a visual stimulus at the preferred orientation combined with optogenetic activation of the orthogonal orientation columns (Fig. 3A). In Fig. 3, all response intensities and contrasts are shown as individual points, and responses for specific cells are examined in the next figures. Once again, we varied the relative drive of these stimuli, by varying contrast in the case of the visual stimulus and by varying optogenetic light intensity in the case of the optogenetic stimulus. Finally, we examined the responses to optogenetic stimulation of columns that matched the preferred orientation of the recorded cell, optogenetic stimulation of columns that were orthogonal to the preferred orientation of the recorded cell, and responses to both stimuli paired, at a variety of optogenetic stimulus intensities (Fig. 3B).

Even before considering responses by single cells, there were obvious differences in the interactions among the different stimulation conditions. Responses of all recorded cells (N=23) to all contrast combinations are shown in Fig. 3C-E. Cross-orientation visual stimulation showed substantial non-linear summation at all contrast levels and response intensities (Fig. 3C), while visual-optogenetic or optogenetic-optogenetic stimuli combined linearly at low-to-moderate response levels and then exhibited sublinear summation at high response levels (Fig. 3DE). These results are incompatible with the idea that the 40% normalization that is found in plaid visual stimulation arises primarily via cortical suppression. Response data are available in Table 3-1 and additional plots are in Extended Data Figure 3-1.

The similarity of the plaid vs. linear sum responses across the conditions where the orthogonal columns were activated optogenetically (Fig. 3D, 3E) raises the question as to whether we find any evidence of a contribution of the cortical circuit in those conditions. Is there a difference in cortical integration if the preferred neurons are activated visually or optogenetically?

To address these questions, we fit the data to a normalization equation (5) (Carandini and Heeger, 2011). The equation allows us to fit additive contributions (*p*-variables) and divisive contributions (*q*-variables) for orthogonal visual stimulation, orthogonal optogenetic stimulation, and preferred optogenetic stimulation.

Analysis of single cell responses indicated that normalization was substantially different across these different stimulus conditions. In the all-visual condition, responses were highly sublinear across all response intensities, and orthogonal stimuli tended to be primarily suppressive (*q_OpOrth_* > 0), as shown in the example cells in Fig. 4A,B,C. In the mixed vision-optogenetic cases, the influence of orthogonal optogenetic stimulation was very small (**Fig. 4DEF**). Only 11 of 23 cells (47.8%) exhibited *p_OpOrth_* and *q_OpOrth_* values that differed significantly from 0 in a nested-F test (all cells exhibited overall optogenetic responses that were significant by a nested F-test, see Methods). Similarly, orthogonal optogenetic stimulation had only slight influences on responses driven by stimulation of the preferred orientation columns (**Fig. 4GHI**).

**Figure 4:**
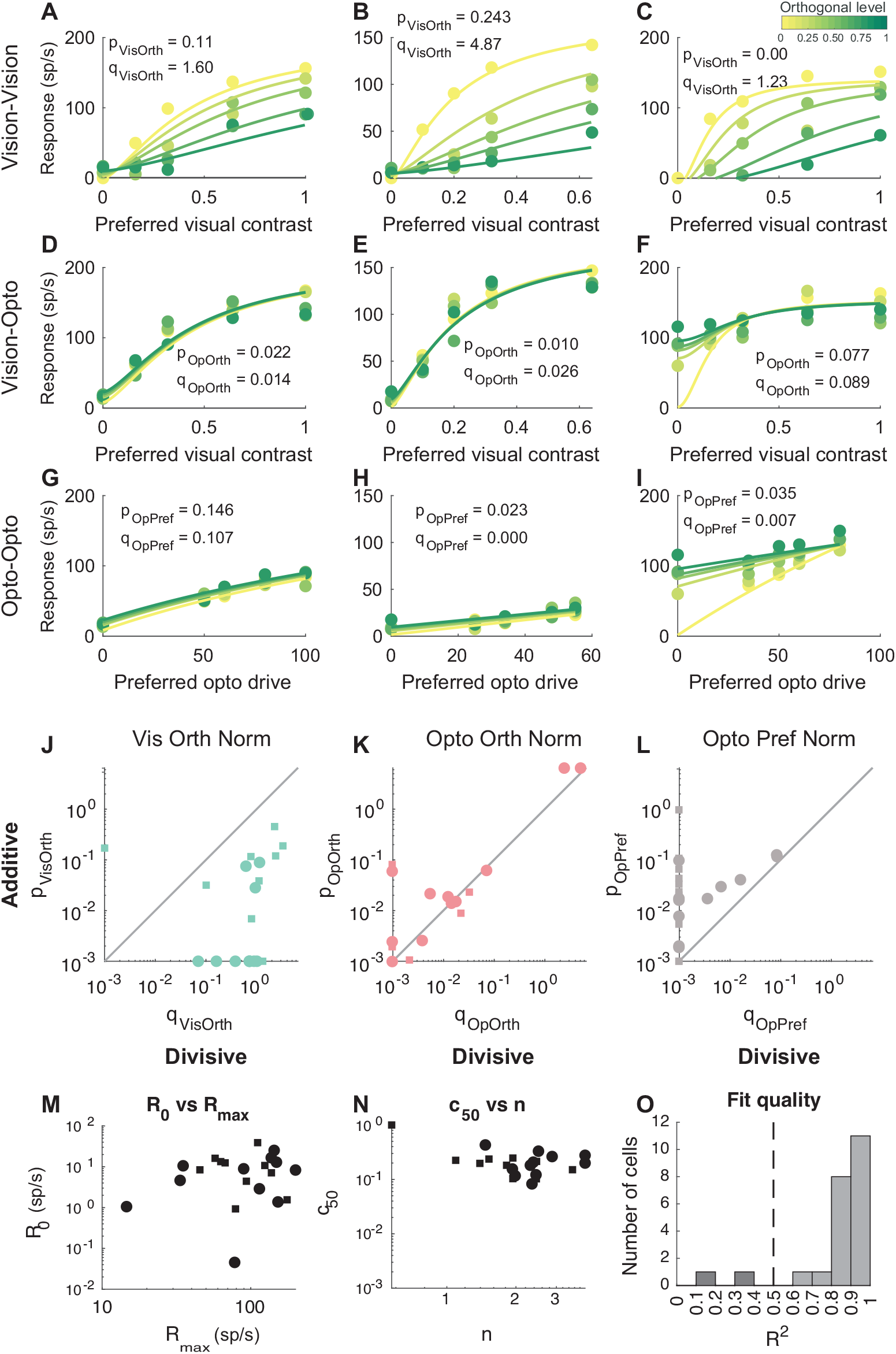
Single unit responses to plaid stimuli. **A**: Response of an example neuron to stimulation at the preferred orientation at different contrasts (X-axis) with five different levels of orthogonal contrast. Orthogonal visual stimulation strongly suppresses response to preferred stimulus. **B,C**: Same for two more cells. **D**: Responses of same cell as A to stimulation at the preferred visual stimulus with simultaneous optogenetic stimulation of orthogonal orientation columns at different drive levels. Stimulation hardly impacts response to the preferred stimulus. **E,F**: Same for neurons in B,C. **G**: Responses when preferred optogenetic columns are stimulated at different levels and orthogonal columns are stimulated optogenetically at different levels. The orthogonal column stimulation has a very small effect. **H,I**: Same for cells in BE,CF. In A,D,G p- and q-based fit parameters are shown. Fit parameters were determined in a two-stage process by fitting purely visual responses first and then optogenetic responses. J,K,L: Comparison of p- (additive contribution) and q- (divisive contribution) parameters for orthogonal visual stimulation **(J)**, orthogonal optogenetic stimulation (K), and optogenetic preferred stimulation (**L**) across all cells. Cells that passed a nested F-test for significant p_OpOrth_ and q_OpOrth_ are shown as circles, while cells without significant p_OpOrth_ and q_OpOrth_ contributions are shown as squares. Orthogonal visual stimulation causes strong divisive normalization. Orthogonal optogenetic stimulation produces much smaller divisive contributions than orthogonal visual stimulation, and the additive components are larger or comparable to the divisive components. Similarly, optogenetic preferred drive is mostly additive. **M,N**: R_0_, R_max_, c_50_, and N for all neurons. **O**: Fit quality histogram (r2) for all cells. 2 cells with r2 < 0.5 excluded.

The overall fit parameters are shown in **Fig. 4JKLMNO**. Divisive factors for orthogonal visual stimulation were on the order of 1 on average with much smaller additive contributions (**Fig. 4J**). By contrast, additive and divisive contributions of orthogonal optogenetic stimulation were quite small, indicating a small contribution to the overall response (**Fig. 4K**). Optogenetic stimulation of preferred columns generally produced an additive response that was almost always weaker than the response to preferred visual stimulation (*p_OpPre f_* < 1) but there was usually a small divisive contribution that was less than or equal to the additive contribution (**Fig. 4L**). The fit quality (*r*^2^) was generally quite good, with all but 2 cells exceeding 0.5 (**Fig. 4O**).

Overall, these data show that optogenetic stimulation of orthogonal orientation columns produced a relatively weak modulation of cortical cell responses, suggesting that orthogonal orientation columns cannot play a major role in cross-orientation suppression responses that are observed with visual stimulation.

### Circuit models that might underlie these responses

The ring model of Rubin et al. (2015) produces strong cross-orientation suppression without direct cross-orientation inhibition (**Fig. 5AB**). Each position in the ring represents a preferred orientation, ranging around the ring from 0^◦^ (= 180^◦^) to 179^◦^, and is modeled by a pair of E and I cells that are reciprocally- and self-connected. Each pair forms a supralinear stabilized network. The synaptic strengths of the horizontal connections across elements of the ring fall off slowly with distance/orientation in a Gaussian manner, and all connection strengths (E-to-E, E-to-I, I-to-E, I-to-I) fall off with the same distance dependence (σ = 32°, as one moves along the ring). The visual input is tuned to stimulus orientation and falls off in a Gaussian manner (σ = 30°) as one moves away from the cells that prefer a given orientation, and optogenetic input falls off similarly but is set slightly broader (σ = 45°). The visual and optogenetic input are provided to both E and I cells. Neurons with different maximum firing rates are simulated by varying the maximum input level provided to different simulations.

**Figure 5:**
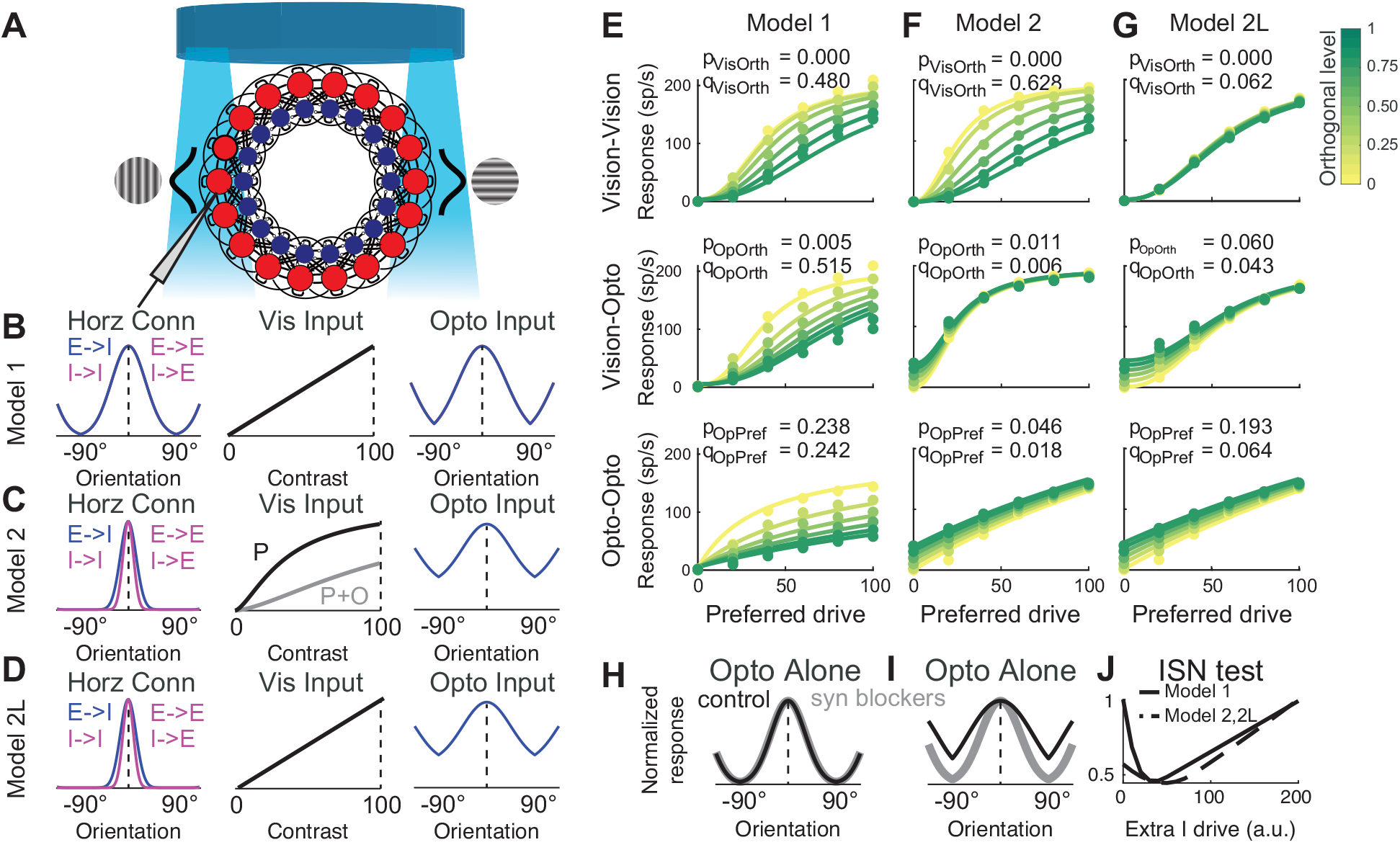
Models of the responses. **A**: Common to all models: Cells are arranged in a ring, with an excitatory (red) and inhibitory (blue) unit at each position. Position corresponds to preferred orientation. The units are coupled in a supralinear stabilized network. Visual and optogenetic inputs are centered at the stimulus orientation, and decrease as a Gaussian function of the difference between the stimulus orientation and the units’ preferred orientation. For visual input, the Gaussian fall off has standard deviation 30°. Visual input strength to E and I cells is identical. The ProjectorScope is depicted as the blue disk above. B: Model 1: The ring model of Rubin et al. (2015). Horizontal connectivity among all cortical cells decreases as a Gaussian of standard deviation 32°. Visual input is linear with contrast. Optogenetic input falls off with width of 45°. C: Model 2: Modified model with less cortical normalization. Now, E to I connections fall off as a Gaussian of 15° width; other connections fall off with 10°. Optogenetic inputs now have a constant pedestal, reflecting the hypothesis that we likely stimulate non-visual neurons that may have broader connectivity. Visual input exhibits cross-orientation normalization. D: Model 2L. Same as Model 2 but with linear visual input, for demonstrating the cortical integration of Model 2. E: Responses of Model 1 to stimulation with combinations of 6 preferred levels of drive and 6 orthogonal levels of drive (same color key as previous figure). Top) preferred and orthogonal are visual; Middle) preferred is visual and orthogonal is optogenetic; Bottom) preferred and orthogonal are optogenetic. Parameters were fit with a two-step procedure and P and Q variables related to visual and optogenetic stimulation are shown. Model 1 shows too much cross-orientation suppression within cortex, indicating that normalization is stronger than we observe in our data. F: Same plots for Model 2. Visual normalization is provided in the visual inputs. Optogenetic normalization within cortex is much weaker and more consistent with the data in 4. G: Same for Model 2L, with linear visual input to showcase the cortical contribution to the cross-orientation normalization of Model 2. H: Normalized response of Model 1 with orientation columns driven optogenetically with and without synaptic blockers. Model 1’s normalized response is identical. I: Normalized response of Model 2 driven optogenetically with and without synaptic blockers. Relative optogenetic tuning width drops with synaptic blockers. J: Tests showing that Models 1, 2, and 2L operate in an inhibition-stabilized network (ISN) regime. Activity of I neuron at preferred orientation as external drive is provided to I cells. Paradoxically, I neuron activity goes down, because the increased inhibition shuts down excitation that was driving the inhibitory neurons, until large external drive dominates. Both models exhibit strong recurrent excitation and inhibition; the difference is the reduction in cross-orientation signals in Model 2.

Simulations of the Rubin et al. model, which we call Model 1, (**Fig. 5E**) exhibit strong cross-orientation suppression by either visual or optogenetic drive. This suppression from orthogonal optogenetic drive is much stronger in this model than we observed in our data.

To capture the responses we observed, we altered the model so that E-to-E, I-to-E, and I-to-I connections were all much more local (10° Gaussian fall-off instead of 32° fall-off), while leaving E-to-I connections at a 15° fall-off to allow much narrower within-cortex normalization (**Fig. 5C**). We called this model Model 2. The E-I cell pairs still exhibited supralinear stabilized network dynamics, but the cross-orientation interactions are much narrower. We dropped the requirement that the cortical circuit itself should provide visual cross-orientation suppression; instead, we provided visual cross-orientation normalization in the feed-forward inputs (Freeman et al., 2002; Li et al., 2006; Priebe and Ferster, 2006; Priebe, 2016) (**Fig. 5C**). Further, we assumed that optogenetic input had a broader impact across orientations to capture our measurements in **Fig. 2CD**, and verified the change in response to stimulation of neighboring orientation columns (mimicking **Fig. 2CD**) in **Fig. 5I**. Responses of this model (**Fig. 5F)** strongly resembled our actual data. Optogenetic activation of orthogonal orientation columns provided only slight modulation of activation of preferred columns, whether by vision or optogenetics.

To demonstrate the within-cortical properties of Model 2, we also ran simulations in a configuration of Model 2 that we call Model 2L where the visual input was completely linear (**Fig. 5D**). As is apparent in simulations (**Fig. 5G**), across-orientation normalization is very weak in cortex in both the visual and optogenetic cases.

Despite strong cross-orientation normalization, both of these Models (1+2) still exhibit inhibition-stabilized dynamics (**Fig. 5J**). If inhibitory neurons are artificially activated, the activity of inhibitory neurons is first reduced, reflecting the removal of excitatory input that was partly responsible for their activity. Only when direct artificial activation becomes very strong does inhibitory activity begin to rise.

### Selective optogenetic stimulation of inhibitory neurons reveals broad classes of functional types

While our results and simulations suggested that within-cortex contributions to cross-orientation suppression might be very small, we wanted to look for some evidence of recurrent activity in cortical networks. We sought to look for direct evidence of the inhibition-stabilized dynamics that the model posits, as has been found in the mouse (Sato et al., 2016; Adesnik, 2017; Sanzeni et al., 2020) and for which evidence has been reported for surround suppression in cat (Ozeki et al., 2009) and ferret (Rubin et al., 2015). A key prediction of ISN-type dynamics is the presence of so-called “paradoxical responses” (Ozeki et al., 2009). If network activity is driven strongly by recurrent connections among excitatory and inhibitory cells, then local excitatory cells are responsible for a significant portion of the synaptic drive to inhibitory cells. If one were to selectively deliver an increase in drive to inhibitory neurons, then excitatory neurons would slow down, but this reduction in excitatory drive would in turn reduce the activity of cortical interneurons. Hence, in the model, the inhibitory cells “paradoxically” respond to small increases in activation with overall reductions in activity (as compared to the increase in activity that might be naively expected).

This prediction is illustrated by simulations in **Fig. 6**. Under a parameter regime where excitatory recurrent connections are set so strongly that the network activity would blow up without inhibition, and inhibitory synaptic strengths are set to be high enough to stabilize this activity (ISN regime), an external increase in drive to inhibitory neurons results in the described paradoxical decrease in activity in interneurons, until the external optogenetic drive to the inhibitory interneurons becomes so strong as to dominate the interneuron responses (**Fig. 6ABC**, same parameters as a single column of the rings of Rubin et al. 2015 and Model 2). On the other hand, if recurrent connections among excitatory and inhibitory neurons are weak (non-ISN), then external drive to inhibitory interneurons merely serves to monotonically increase the activity of these interneurons (**Fig 6DEF**).

**Figure 6:**
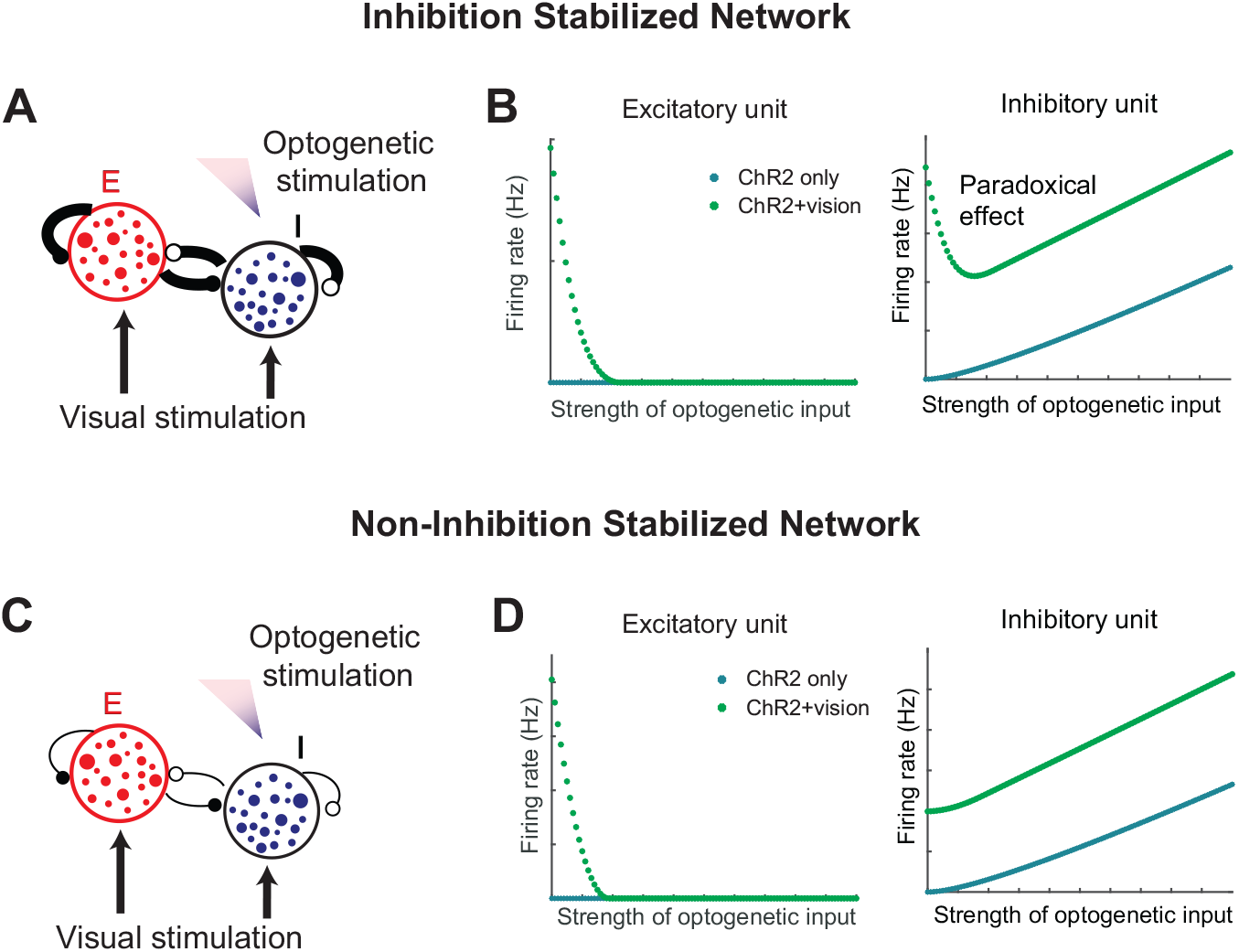
Differential responses to optogenetic interneuron stimulation in ISN and non-ISN models. **A**: In the simulation of ISN ring model (illustrated here as one pair of E and I), the E and the I units have strong recurrent connections. Visual stimulation provides inputs to both the E and the I units, and optogenetic stimulation provides inputs to the I units only. **B** (left): Excitatory unit responses in the ISN. When the inhibitory units are activated by ChR2 stimulation, the excitatory units have no responses. When visual stimulation is provided together with ChR2 stimulation, the excitatory units respond strongly to visual stimulation when the inhibition is weak and reduce firing rates as the inhibition becomes stronger. B (right): Inhibitory unit responses in the ISN. When the inhibitory units are activated by ChR2 stimulation but are not otherwise active in the circuit, they monotonically increase firing rates as the stimulation becomes stronger. When ChR2 stimulation is provided in the presence of visual stimulation, the firing rates decrease before increasing, creating a “dip” shape in the response curve that is characteristic of the ISN. **C**: In the simulation of the non-ISN model, the ring structure is maintained as in the ISN but the recurrent connections are weak. **D** (left): Excitatory unit responses in the non-ISN network. Activation of the interneurons suppresses the excitatory unit activity. D (right): Inhibitory unit responses in the non-ISN network. In the non-ISN network, the inhibitory units show direct responses to light in a monotonically increasing manner. When ChR2 stimulation is combined with visual stimulation, the inhibitory units also respond in a monotonically increasing manner and the “dip” is absent.

To test these predictions, we prepared ferrets with a virus (AAV9-mDlx-ChR2-mCherry-Fishell-3) that restricts the expression of channelrhodopsin-2 to inhibitory neurons (Dimidschstein et al., 2016). We delivered wide-field white light to stimulate the brain surface, and the light intensity was modulated to achieve different levels of external drive to inhibitory neurons. Optogenetic stimulation was presented with and without visual stimulation at the preferred orientation with different contrasts, in order to understand how the activated cortex would be modulated by the external optogenetic increases in inhibitory drive. In these experiments, a 32-channel probe (Plexon S-probe) was used to achieve better yield of recording both excitatory and inhibitory cells.

We observed a variety of response profiles to combined visual stimulation and optogenetic stimulation of interneurons. Some cells exhibited no response to optogenetic interneuron stimulation alone, but exhibited strong suppression of visual responses when optogenetic drive was strong. We labeled these neurons as putative excitatory neurons (**Fig. 7A**, bottom row).

**Figure 7:**
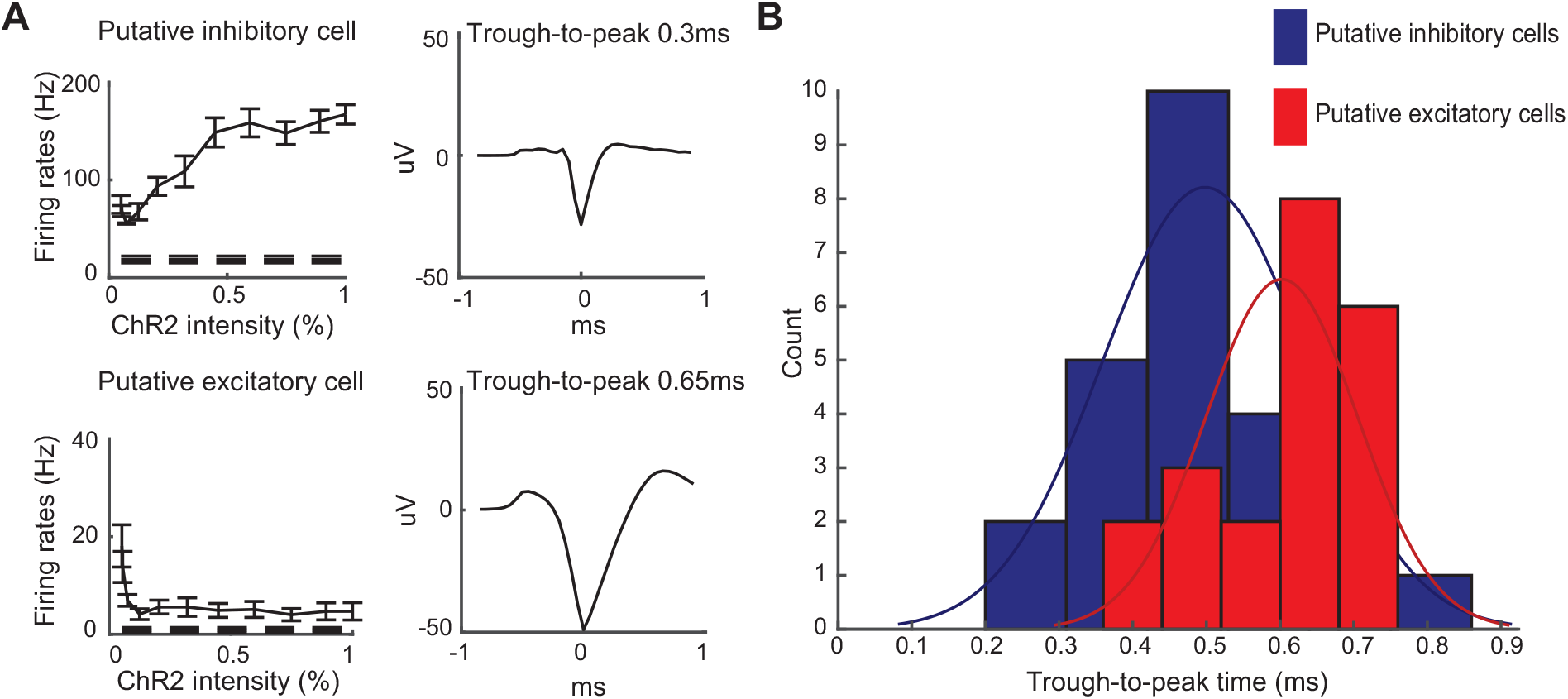
Optogenetic stimulation of ferrets expressing mDlx-ChR2 revealed neurons with putative excitatory and inhibitory signatures. **A**. Left) Responses of an example putative inhibitory cell (top) that shows direct activation by light stimulation, and a putative excitatory cell (bottom) that is strongly inhibited by light. Raw responses are shown, without visual stimulation, and without any subtraction of background firing rates (dashed line). Right) Zoomed-in views of the average action potential waveform from each neuron along with the trough-to-peak time. **B**. Histograms of trough-to-peak times of putative inhibitory cells and putative excitatory cells. Cells’ identities were first determined by their response profiles to optogenetic stimulation and then the trough-to-peak times were calculated and summarized. Gaussian density fits to the histograms are shown in green. On average, the putative excitatory cells showed longer spike durations than the inhibitory cells, although there were putative inhibitory cells that exhibited wider spikes, consistent with the variety of interneuron types found in cortex (fast-spiking and regular-spiking).

We also observed response profiles that we imagined arose from inhibitory neurons. These cells, labeled as putative inhibitory neurons, exhibited strong responses to strong optogenetic stimulation when it was presented either with or without visual stimulation (**Fig. 7A**, top row). The putative inhibitory neurons always responded directly to light and, on average but not always, exhibited shorter spike duration than the putative excitatory cells (**Fig. 7B**).

The putative inhibitory interneurons were not uniform in response profile, but instead exhibited a range of responses. At one extreme, responses from some putative inhibitory interneurons resembled the responses from interneurons in an ISN-like circuit (**Fig. 8A**, top row). During high contrast visual stimulation, the activity of these interneurons was suppressed for weak optogenetic activation, consistent with the idea that increased drive to interneurons was reducing excitatory activity, which in turn decreased recurrent inhibitory activity. Without simultaneous visual stimulation, these were less clear, given the low spontaneous firing rates of most neurons. At another extreme, we also observed several neurons that exhibited monotonic increases in responses to optogenetic drive (**Fig. 9A**, middle row). These results suggest that there are multiple ways in which interneurons can be interconnected with cortical circuits. The full range of tuning profiles that we observed is projected onto its first two principal components and plotted in **Fig. 8B**. Within the putative inhibitory population, the data are more consistent with a continuum rather than discrete clusters.

**Figure 8:**
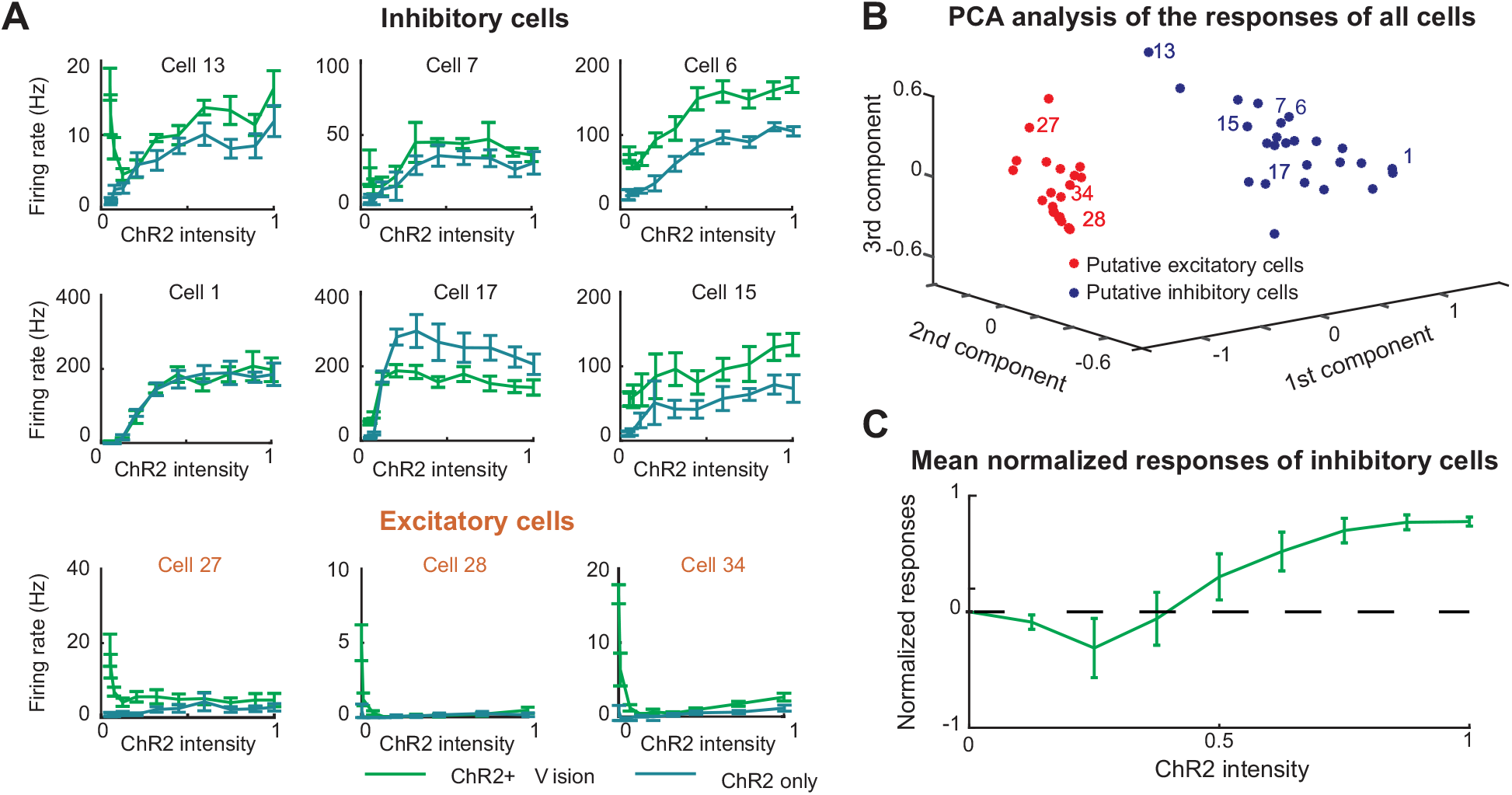
The influence of direct interneuron stimulation on visual responses in putative excitatory and inhibitory interneurons. **A:** Representative unit responses from the optogenetic inhibition experiments. Raw mean firing rates (without any background rate subtraction) are shown for increasing optogenetic stimulation alone (blue) or in the presence of a high contrast visual stimulus (green). Top) Putative inhibitory neurons that responded like neurons in the ISN simulation. Middle) Putative inhibitory neurons that did not respond like the ISN simulation. Instead, the firing rates either saturated or increased at low light intensities and then decreased at high light intensities. Bottom) Putative excitatory neurons. B: Projection of response profiles to visual stimulation and increasing optogenetic interneuron activation onto the first 3 principal components. Putative excitatory and inhibitory neurons are well separated, but there is diversity of responses within the putative inhibitory population. Numbers indicate the example neurons displayed in A. C: The grand average of the normalized ChR2+Vision responses of all putative inhibitory cells. For every inhibitory cell, the responses were normalized in such way that the average response at 0 light intensity is normalized to 0 and the average maximum response is 1.

**Figure 9:**
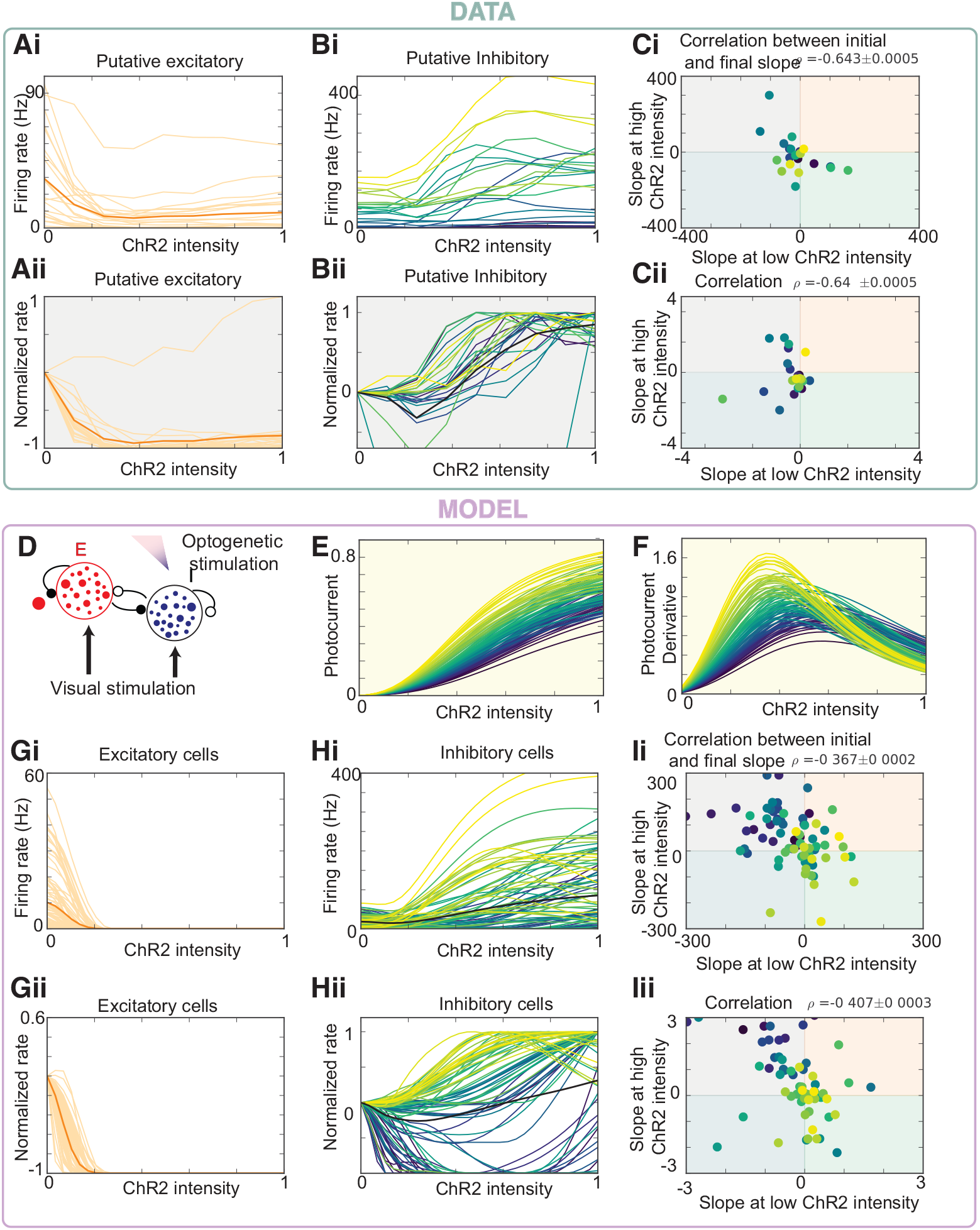
A network model with heterogeneous connectivity and opsin activation reproduces the diversity of interneuron responses, including paradoxical suppression. TOP: Ai: Putative excitatory cells (as defined in Fig. 7) as a function of the intensity of the optogenetic input to the inhibitory cells. Aii: Same as Ai but normalized such that the curves start from zero and were maximal at 1 (as in Fig. 8c). Bi: and Bii: Putative inhibitory cells. The paradoxical effect (i.e. that some cells decrease their activity upon optogenetic drive) is better revealed by normalizing the cells. Ci: The slope of each cells response to a weak laser input is negatively correlated with the slope at a large laser input. In other words, the cells that did not respond paradoxically tended to saturate at smaller laser intensities. Cii: Negative correlation for normalized responses. BOTTOM: D: Heterogeneous model with N neurons (N=500 in simulations). E: Model of optogenetic current for each cell as a function of laser intensity as given by Eq. 12. F: Derivative of the photocurrent. Gi: Excitatory cells as a function of the intensity of the optogenetic input to the inhibitory cells. Gii: Same as Ai but normalized such that the curves start form zero and are maximal at 1. Hi: and Hii: Inhibitory cells. The paradoxical effect in the model is better revealed by normalizing the cells, as in the data. Ii: and Iii: The model captures the negative correlation between the slope in response to a weak laser and the slope at a large laser input for both normalized and non-normalized responses.

The diversity of responses from putative interneurons raises the question as to the nature of the overall impact of cortical interneurons on the circuit. Of course, interneuron connectivity could be very specific, but as a point of interest we calculated the grand average of the normalized responses of all the inhibitory cells to combined optogenetic stimulation and visual stimulation (**Fig. 8C**). This grand average would reflect the inhibitory influence on the cortical network if interneuron types were pooled unselectively. The grand average tuning curve exhibits an empirical dip below zero for weak external input, although no point on the curve is significantly below 0 with a p-value of less than 0.05.

Diversity in interneuron responses to optogenetic stimulation could arise from diversity in the response of different inhibitory cell types, due to heterogeneity in neural connections, or due to the heterogeneity in the opsin expression across inhibitory cells. We created a large-scale EI network model with heterogeneous, randomly distributed connectivity (Fig. 9). We modeled the photocurrent to each cell as a Hill equation as reported experimentally (Asrican et. al. 2018), and the activation of each interneuron to vary with depth, as expected from scattering of light across the cortical tissue (Yona et. al. 2018). In this model, we observed a heterogeneous range of response profiles in optogenetically-activated interneurons, as in the data. The model not only captures the initial response of I cells to laser power, but also the negative correlation between the effect of the laser at low intensity and the effect of the laser at high intensity to each cell: If the cell is initially excited by the laser, then it tends to quickly saturate for large laser power, whereas if it responds paradoxically to the laser at low power (i.e. if it has a negative response to weak excitatory input) then its response does not saturate at large laser power. These results suggest that diversity in such properties as synaptic connectivity, optogenetic activation strengths, and cellular thresholds could underlie the variable response profiles we observed in **Figure 8**.

## 5. Discussion

We performed combined visual and patterned optogenetic stimulation to test how cortical circuits responded to multiple inputs. We found that cortical stimulation that was optically restricted to specific orientation columns caused activity that spread non-selectively to neighboring columns. We used this protocol to examine the cortical contributions to contextual modulation by stimulating different cortical columns visually and optogenetically. We found evidence for cross-column cortical normalization, but much less than would be needed for a purely cortical mechanism to account for cross-orientation suppression. We found a wide range of interneuron couplings to the circuit, including those that responded to weak optogenetic activation by reducing their activity, consistent with inhibition stabilized networks.

### Nonspecific spread of activity with respect to orientation columns

We found that column-based patterned optogenetic stimulation did not respect the boundaries of orientation columns, although local activation was typically restricted around the recorded neuron. While it is difficult to exclude the possibility that this unselective spreading is due in part to activation of passing axons and dendrites of neurons in other columns, blocking NMDA and AMPA receptors revealed that the influence of horizontal connections played a large role. Viruses that cause strong expression of ChR2 that is targeted to the soma could help manage concerns about passing dendrites, but the one soma-targeting virus we were able to try did not cause sufficient expression to produce strong responses in vivo with our stimulation system (Baker et al., 2016). Our results are consistent with a study that performed sparse stimulation of nearby orientation columns in tree shrew (Huang et al., 2014). One might have expected to find species differences between tree shrew and ferret because the tree shrew primarily exhibits length-summation in layer 2/3 (Chisum et al., 2003) while ferret exhibits both length-summation and surround suppression (Rubin et al., 2015; Popovic et al., 2018), but both species showed non-specific spread of activity in layer 2/3.

An important corollary of nonspecific spread is that column-level stimulation of cortical neurons is unlikely to provide an optimal stand-in for visual activation, e.g. in a visual prosthesis. Visual stimulation only sparsely activates cortical neurons (Rochefort et al., 2009; Haider et al., 2010; Smith et al., 2015), and there appear to be cells that are not activated by visual stimulation or are only activated in conjunction with some non-visual stimulus (Saleem et al., 2018), whereas an optogenetic stimulus will activate all cells in a column. It is possible that expression of optogenetic channels restricted to LGN axons may allow more specific stimulation of visually-driven neurons in particular.

### Linear vs. non-linear interactions of visual and optogenetic signals

Huang and colleagues (2014) used AAV viruses to cause broad expression of channelrhodopsin in excitatory neurons in the tree shrew. Optogenetic activation using small spots of light targeted to preferred or orthogonal columns added linearly over a wide range of firing rates. Further, visual stimulation with the preferred orientation combined with optogenetic stimulation of preferred columns also showed linear summation.

In macaque, Nassi et al. (2015) used lentivirus to cause more localized expression of C1V1 in excitatory neurons of the macaque. These investigators stimulated broadly with an optical fiber and observed a variety of facilitatory and suppressive interactions to joint visual stimulation and optogenetic stimulation that was not specific to particular columns of the orientation map. The vast majority of these neurons exhibited sublinear summation of visual and optogenetic signals.

Histed (2018) used transgenic approaches to cause expression of ChR2 in mouse visual cortical neurons, and found that visual and optogenetic inputs summed in a largely linear manner, though with sublinear summation at higher firing rates. Another study in the mouse, which used optogenetic antidromic activation of callosal inputs, found that callosal inputs facilitated responses at low visual contrasts but suppressed responses at higher visual contrasts (Sato et al., 2014).

Our data (**Fig. 3**) also shows evidence of nearly linear summation when neurons exhibited low firing rates, which became sublinear as neurons exhibited larger firing rates.

### In layer 2/3, cross-column suppression is insuffcient to account for cross-orientation suppression to visual stimuli

Cortical neurons exhibit weaker responses to a preferred oriented stimulus when the stimulus is combined with an orthogonally-oriented stimulus (Bishop et al., 1973; Morrone et al., 1982; Bonds, 1989; DeAngelis et al., 1992; Cavanaugh et al., 2002; Smith et al., 2006; Busse et al., 2009; MacEvoy et al., 2009). Experiments in the last 20 years have shown that direct cross-column inhibition is unlikely to underlie this phenomenon, as synaptic conductance measurements of both excitatory and inhibitory inputs peak at the preferred orientation and are relatively weak at the orthogonal orientation (Anderson et al., 2000), and both excitation and inhibition are reduced by the addition of an orthogonal grating stimulus to a preferred-orientation grating stimulus (Priebe and Ferster, 2006).

A more recent model suggested that nonlinear circuit properties induced by supralinear single-cell input/output functions could explain the change from cross-orientation facilitation for weak stimuli to cross-orientation suppression for stronger stimuli. Furthermore, because the network was inhibition-stabilized for stronger stimuli, the model could explain cross-orientation suppression with a combined reduction of excitatory and inhibitory conductances (Ozeki et al., 2009; Rubin et al., 2015; Anderson et al., 2000; Priebe and Ferster, 2006).

Alternatively, other past models suggested that there was no need for any cortical explanation of cross-orientation suppression. These models suggest that cross-orientation suppression can be largely accounted for by changes in LGN inputs to V1 cells along with V1 spike threshold (Lauritzen et al., 2001; Freeman et al., 2002; Li et al., 2006; Priebe and Ferster, 2006; Priebe, 2016). Recently, this idea was tested directly by silencing cortex with optogenetic stimulation of parvalbumin neurons in mice, and normalization in feed-forward input (as viewed in the membrane potential) was unaltered with cortical suppression.

Our optogenetic stimulation results were inconsistent with a cortical explanation for cross-orientation suppression. Under the hypothesis that optogenetic stimulation is in any way like a visual stimulation, stimulation of orthogonal columns should have produced a strong suppression in the orthogonal columns. Instead, we observed a moderate spreading of cortical responses. Combined visual and optogenetic stimulation produced responses very similar to the linear sum of the individual stimuli for moderate response strengths, while paired stimulation of visual stimuli of moderate contrast produced strongly sublinear responses. In our model that best matched the data, there was some weak cross-column suppression, but the cross-column suppression was much smaller than is required to produce the cross-orientation suppression seen using visual stimuli.

Now experiments have shown that feed-forward information should (in principle) be able to provide cross-orientation suppression (Freeman et al., 2002; Li et al., 2006; Priebe and Ferster, 2006), silencing cortical activity does not alter subthreshold cross-orientation suppression (Barbera 2022), and activating neighboring columns does not suppress cross-orientation suppression (this paper). On the basis of all of this evidence, it is highly likely that cross-orientation suppression is feed-forward in nature.

### Inhibition-stabilized dynamics and paradoxical responses

Another goal was to examine whether we could find direct evidence of inhibition-stabilized dynamics (Suarez et al., 1995; Tsodyks et al., 1997; Ozeki et al., 2009; Rubin et al., 2015) in ferret visual cortex. Inhibition-stabilized dynamics have been observed in surround suppression in cat visual cortex (Ozeki et al., 2009) and mouse visual cortex (Sato et al., 2016; Adesnik, 2017; Sanzeni et al., 2020), mouse somatosensory cortex (Sanzeni et al., 2020), and mouse motor cortex (Sanzeni et al., 2020). Sanzeni et al (2020) also looked for paradoxical responses by direct optogenetic stimulation of interneurons, and found that paradoxical responses in awake animals could be evoked by stimulating either all interneurons or just PV neurons when they were targeted in a transgenic manner. Interneuron receptive field properties differ between species that have columnar features beyond retinotopic maps and those that only have retinotopic maps; for example, interneurons in cat and ferret visual cortex can be highly tuned for orientation (Hirsch et al., 2003; Cardin et al., 2007; Wilson et al., 2017), while a majority of interneurons in mouse cortex are not (Sohya et al., 2007; Niell and Stryker, 2008; Liu et al., 2009; Kerlin et al., 2010). Here we showed that a subset of ferret interneurons also exhibited characteristic paradoxical behavior with direct stimulation.

We observed heterogeneous interneuron responses to increasing optogenetic stimulation. These responses resemble the heterogenous paradoxical/non-paradoxical responses that Sanzeni et al. (2020) and Mahrach et al (2020) observed in experiments using viral-mediated infection of PV+ neurons. In ferret, we used a viral promotor that caused expression of ChR2 in a wide variety of interneuron classes (Dimidschstein et al., 2016), but interneuron subclass-specific viruses are now becoming available for non-rodents (Mehta et al., 2019; Vormstein-Schneider et al., 2020). Future experiments will be needed to determine if some of the heterogeneity we observed could be due to different interneuron subtypes.

## Acknowledgements

This work was funded by NIH EY029999 (KDM + SDV), NSF 1707398 (KDM + AP), Gatsby Charitable Foundation GAT3708 (KDM + AP), and Swartz Foundation (AP). We thank members of Miller lab and Van Hooser lab for comments on the work. We thank David Fitzpatrick’s lab for providing AAV9-mDlx-ChR2-mCherry-Fishell-3.

## 6. Contributions

KDM and SDV designed experiments; SW designed ProjectorScope 2.0 and carried out experiments, modeling, and analysis; AP provided models and interpretations. SW and SDV wrote the paper with input from AP and KDM.

